# Genome evolution in parthenogenetic nematodes shaped by chromosome rearrangements and rare sex

**DOI:** 10.1101/2025.04.24.650375

**Authors:** George Chung, Lewis Stevens, Xiaoguang Dai, Scott Hickey, Karin Kiontke, Priyesh Rughani, Manuela R. Kieninger, Joanna C. Collins, Sissel Juul, Fabio Piano, David H. A. Fitch, Mark Blaxter, Kristin C. Gunsalus

## Abstract

In contrast to their dioecious relatives, members of the parthenogenetic *Diploscapter* nematode genus harbour their entire genome within a single pair of highly heterozygous chromosomes. To examine how this unusual karyotype relates to the evolution of parthenogenesis, we generated chromosome-level assemblies for two species in this clade: *Diploscapter pachys* and *Diploscapter coronatus*. Sequence comparisons revealed that the two genomes are colinear across their entirety, and that multiple ancestral chromosome fusions and extensive genomic rearrangements preceded the divergence of these two species. The presence of shortened telomeres and extended subtelomeric repeats suggests that the fusions arose from defects in telomere function in the lineage. Our analysis also identified an introgression event after divergence of the two species, suggesting that their parthenogenetic lifestyle may have been punctuated by rare sexual reproduction. These findings shed new light on how telomere loss, chromatin architecture, and reproductive strategies interconnect in shaping chromosome evolution.

---

Parthenogenesis, or reproduction without fertilization, has evolved independently multiple times in metazoans, indicating that an asexual reproductive mode can emerge in different genetic or genomic contexts. Parthenogenic animals have arisen from interspecific hybridization^1,2^, ploidy changes^3,4^, or both^5–7^. Oocytes capable of embryonic development in the absence of fertilization may arise either by skipping the reductional meiotic division^8^ or through the fusion of two haploid meiotic products^9^. Over time, individual haplotypes in a diploid ameiotic parthenogen accumulate mutations independently, leading to their sequence divergence – a phenomenon known as the Meselson effect^10,11^.

Like other ameiotic and parthenogenetic organisms, diploid nematodes in the *Diploscapter* genus have highly heterozygous genomes^8,12^. However, the genome of *D. pachys* maintains some homozygosity, suggesting that some form of recombination or gene conversion may still be still possible. Parthenogenetic *Diploscapter* nematodes are also unusual in Metazoa in having a single pair of chromosomes^8,12^ resulting from ancestral chromosome fusion events^8^.

The published *D. pachys* and *D. coronatus* genome assemblies, generated primarily from short reads, were fragmented (∼500 to ∼10,000 scaffolds) and lacked identifiable telomeric ends^8,12^. The order of fusions that generated the *D. pachys* single chromosome from the ancestral linkage groups of rhabditid nematodes (Nigon elements^13^) was not known, but most scaffolds could be mapped to Nigon elements.

These assemblies therefore left open several important questions about how the *Diploscapter* genome structures relate to their evolution and the parthenogenetic mode of reproduction. Are the *Diploscapter* chromosomes linear and capped by telomeres, or circular? If they are linear, are the two homologous chromosomes within one *Diploscapter* species colinear throughout their entire lengths, or do they contain rearrangements incompatible with meiotic recombination? What was the identity of the ancestral chromosomes prior to the fusion events, and are the fusion events causally linked to the absence of telomeres, as hypothesized previously^8^? Without a complete haplotypic chromosomal assembly, we could not determine if the homozygous region in *D. pachys* is fully continuous or comprises many islands of homozygosity, which could distinguish between a single ancestral homogenizing event or ongoing processes that would require active recombination machinery.

To fully define the overall genome structure, we employed two long-read sequencing technologies aided by chromatin conformation capture information to generate haplotype-resolved, chromosomal-level assemblies of the *D. pachys* and *D. coronatus* genomes. Our analyses show that the chromosomes in the two *Diploscapter* species are linear and capped by arrays of short telomeric repeats and long subtelomeric repeats. Despite considerable nucleotide divergence within and between the two species, the chromosomes are largely syntenic. One 18-Mb terminal region of the *D. pachys* genome is identical between two haplotypes, indicating that mechanisms capable of homogenization between chromosomes were present in the recent past, although no meiotic recombination has been detected in either species under laboratory conditions. Sequence comparisons also provide evidence of recent introgression of genetic material from one *Diploscapter* species to another, indicating that reproduction involving males was at one point in time possible between them.

## RESULTS

### Chromosome-level genome assemblies for *D. pachys* and *D. coronatus*

To understand the overall genomic structures of *D. pachys* and *D. coronatus,* we used two long-read sequencing methods to assemble the two genomes, complemented by chromatin contact maps to scaffold the assembled contigs (**Supp. Table 1**). *D. coronatus* was sequenced using PacBio HiFi technology (Pacific Biosciences, Menlo Park, USA), while Nanopore technology (Oxford Nanopore Technologies, Oxford, UK) was used to sequence the *D. pachys* genome. Scaffolding using chromatin contact data (HiC or PoreC) revealed that the nuclear genomes each contain two homologous chromosomes between 80-90 Mb long (**Fig. 1a-b**). The chromatin contact maps also revealed that the chromosomes were linear, with distinct ends (**Fig. 1c-d**). In both species, the homologous chromosomes were highly heterozygous. We named the shorter of the pair homolog A and the longer one homolog B.

**Fig. 1:**
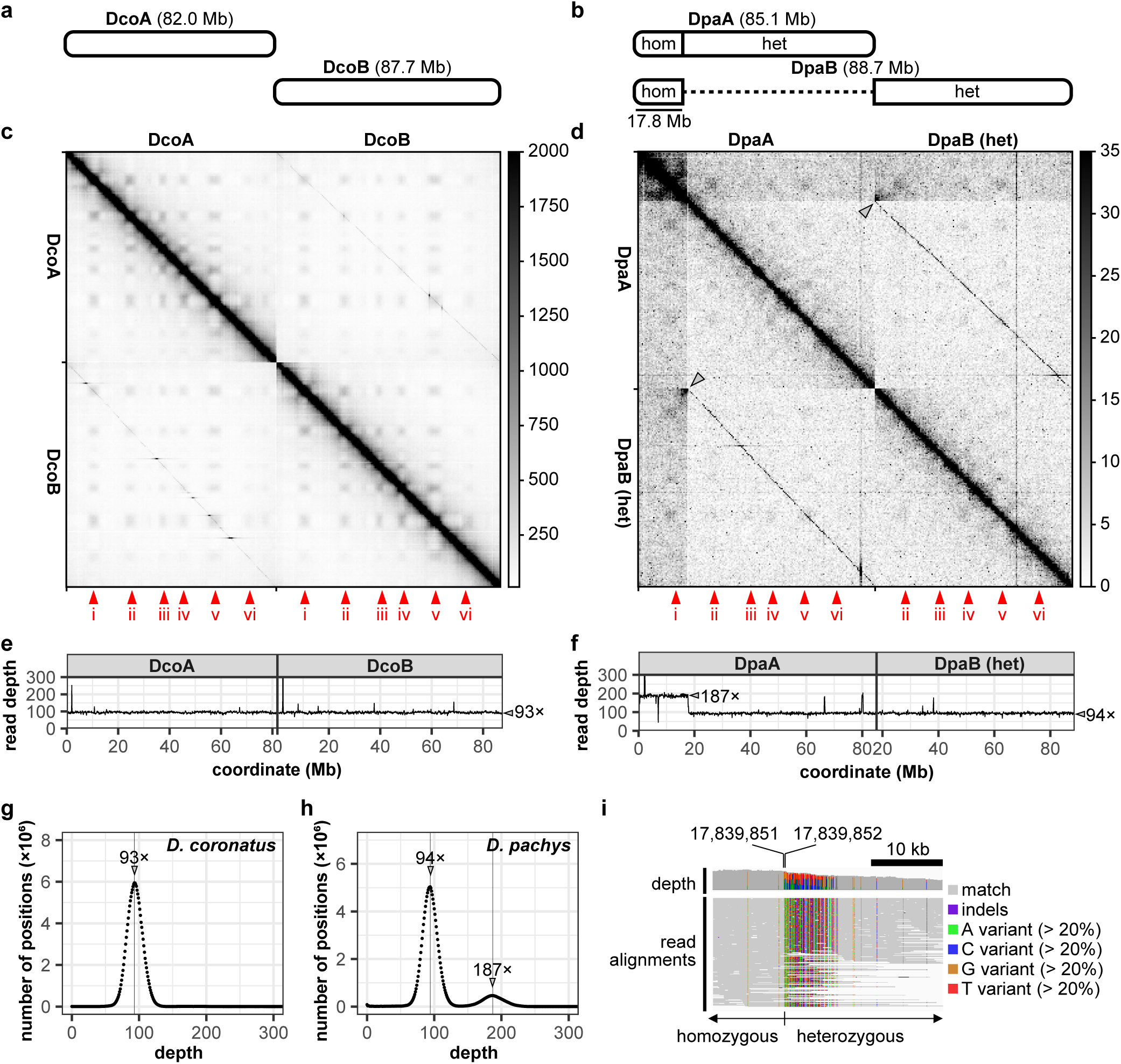
The genome assemblies of *D. coronatus* and *D. pachys*. **a-b,** Schematic of the two nuclear chromosomes in *D. coronatus* (**a**) and in *D. pachys* (**b**). **c-d,** Hi-C matrix plot for *D. coronatus* (**c**) and PoreC matrix plot for *D. pachys* (**d**), both at 250,000-bp resolution and revealing areas of high intra– and inter-chromosomal contact, labelled i to vi (red arrowheads). The homozygous region in the *D. pachys* assembly is connected to both DpaA and DpaB (**d,** grey arrowheads). **e-f,** Read coverage over *D. coronatus* (**e**) and *D. pachys* (**f**) chromosome lengths. *D. pachys* possesses a homozygous region at the left end of DpaA. **g-h,** The number of base pairs with a specific read depth in *D. coronatus* (**g**) and in *D. pachys* (**h**). **i,** The *D. pachys* homozygous-heterozygous junction on homologue A – between nucleotide 17,839,851 and 17,839,852 – is revealed by chromosome B reads mis-mapping to chromosome A. Visualization as displayed on Integrative Genomics Viewer^54^.

In *D. coronatus*, the heterozygosity spans the entire lengths of both chromosomes, supported by ∼93× read coverage for each haplotype (**Fig. 1e, g**). In *D. pachys*, we identified a continuous stretch of homozygosity (17.8 Mbp) at one end of both homologous chromosomes (henceforth referred to as the “left” end). This homozygous region is supported by three observations. First, as in our previous assembly^8^, its read coverage is twice the global average (**Fig. 1f, h**, ∼187×). Second, the *D. pachys* chromatin contact map reveals that this region is represented once in the assembly and is physically connected to two distinct sites on each chromosome (**Fig. 1d**, grey arrowheads). Third, sequence alignments of reads spanning the junction of homozygosity and heterozygosity reveal a clear pattern of mismatches due to the erroneous mapping of homolog B reads onto homolog A (**Fig. 1i**). The continuity of homozygosity across the entire region suggests that it arose from a single recent homogenization event rather than a series of small, segmental gene conversion events. Finally, we did not detect any evidence for programmed DNA elimination, a common trait in many rhabditid nematodes^13–15^.

### Each *Diploscapter* chromosome end is capped by long subtelomeric and short telomeric repeat arrays

While the telomeres of *Diploscapter* chromosomes were not identified previously^8,16^, chromatin contact maps clearly show the presence of distinct chromosome ends (**Fig. 1c-d**). All four chromosome assemblies contain terminal arrays of ∼3-kb tandem repeats, each showing similar but nonidentical repeat motifs. Read depths for these repetitive arrays are much higher than the genome average for all but the left (homozygous) end of *D. pachys* chromosomes (**Fig. 2a-b**), suggesting that they are much longer than they appear and are thus not fully resolved in the assembly (**Supp. table 2**).

**Fig. 2:**
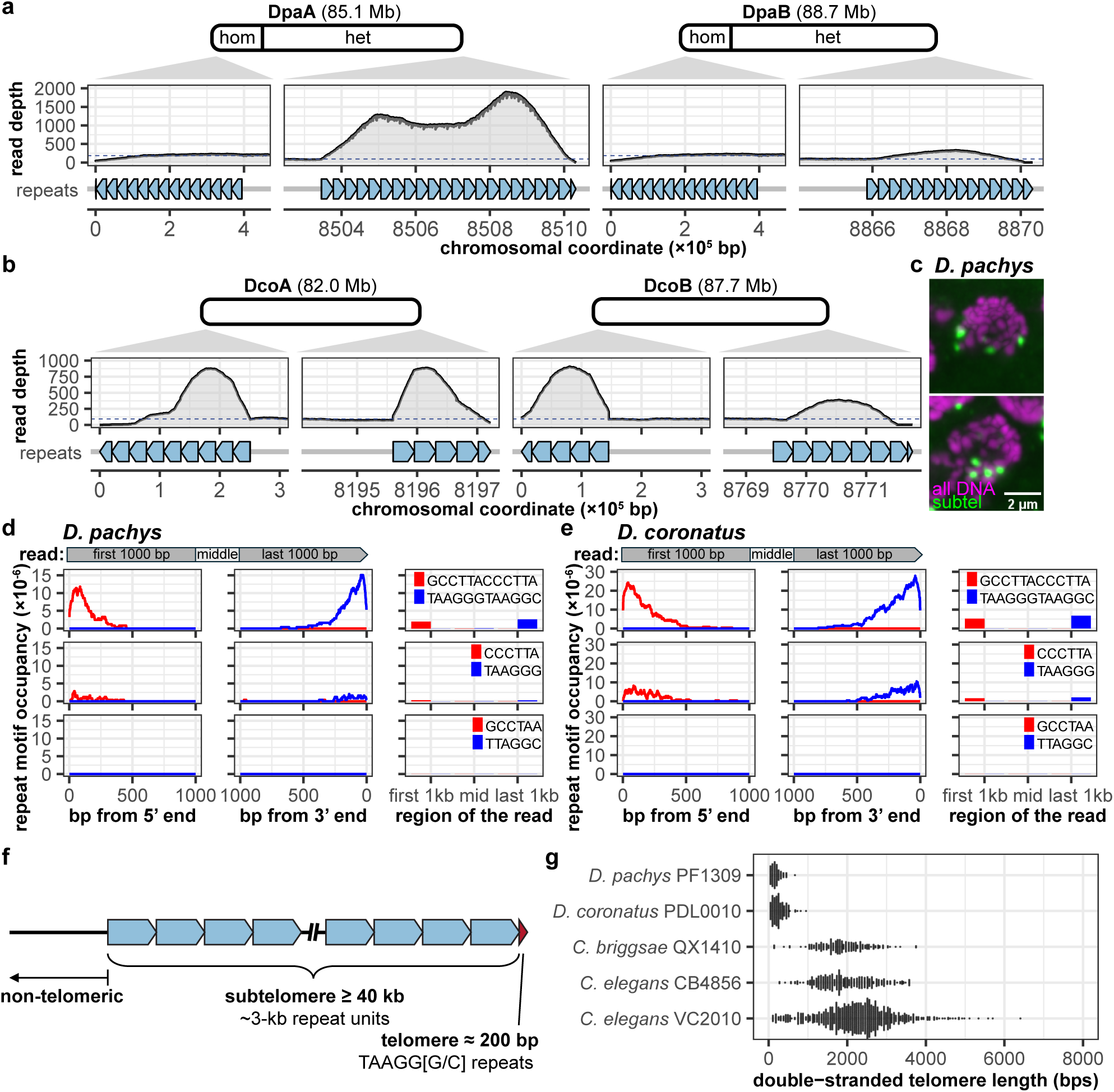
*Diploscapter* has long repetitive subtelomeres and short telomeres. **a-b,** The read coverage is plotted on top, followed by a schematic of the subtelomeric repeat structure, and the chromosomal assembly coordinates. **a,** The ∼3-kb subtelomeric repeats at the ends of *D. pachys* chromosomes. “hom” = homozygous region between DpaA and DpaB. **b,** The ∼3-kb subtelomeric repeats at the ends of *D. coronatus* chromosomes. **c,** FISH staining of *D. pachys* oocyte nuclei using the *D. pachys* ∼3-kb repeat sequence as the probe reveals that the ∼3-kb repeats are near the chromosome ends. **d,** Occupancy of tandem repeat motifs TAAGGGTAAGGC (top), TAAGGG (middle) and TTAGGC (bottom) at the ends of *D. pachys* sequencing reads. Mirrored pattern of the reverse complement motif (red) suggests the original (blue) is a bona fide telomeric motif. **e,** Occupancy of tandem repeat motifs TAAGGGTAAGGC (top), TAAGGG (middle) and TTAGGC (bottom) at the ends of *D. coronatus* sequencing reads. The canonical rhabditid motif TTAGGC is not represented in either species. **f,** Schematic representation of the *Diploscapter* chromosome end, inferred from the sequencing reads and TeloSearchLR results. **g,** Telomere length estimates from long sequencing reads. *Diploscapter* telomeres are short compared to telomeres from nematode relatives from *Caenorhabditis.* Data are derived from this work and long-read sequencing libraries from *C. briggsae* QX1410^41^, *C. elegans* CB4856 (“Hawaiian”)^93^, and *C. elegans* VC2010 (“N2 reference”)^92^.

Although FISH signals from these arrays localize to *D. pachys* chromosome ends in micrographs (**Fig. 2c**), most rhabditid nematode genomes contain telomeres with a canonical 6-bp motif, TTAGGC^17,18^. This led us to wonder whether these long tandem arrays represent true *Diploscapter* telomere sequences or instead may be subtelomeric repeats. To address this question, we used TeloSearchLR^19^ to directly examine long sequence reads for repeat motifs with a strong positional bias. These searches revealed a dominant 12-bp terminal repeat motif in both *D. coronatus* and *D. pachys*, TAAGGGTAAGGC (**Fig. 2d-e**, top), along with a lower-frequency 6-bp motif, TAAGGG (**Fig. 2d-e**, middle) – both of which result from the frequent (but not entirely consistent) alternation of TAAGG<u>G</u> and TAAGG<u>C</u>. These TAAGG[G/C] arrays are located distal to the ∼3-kb tandem repeats in individual reads, indicating that they constitute the true telomeric repeats in these species (**Fig. 2f**). The typical TTAGGC^17,18^ motif is not enriched (**Fig. 2d-e**, bottom), likely due to sequence divergence and possible heterogeneity of the unidentified telomerase RNA template. The telomeric repeat motifs occupy an average of 182 ± 24 bps (*D. pachys*, 95% CI) and 259 ± 25 bp (*D. coronatus,* 95% CI) at the ends of sequencing reads, much shorter than the ∼2 kb telomeric repeat arrays detected in long-read libraries from *Caenorhabditis* relatives (**Fig. 2g**). While these *Diploscapter* telomeres may be maintained by a telomerase ortholog (**Supp. fig. 1**), we could not identify a telomere-binding OB fold protein ortholog (MRT-1, POT-1/2/3 in *C. elegans*) in either of the new *D. pachys* or *D. coronatus* assemblies, consistent with our previous findings for *D. pachys*^8^.

### *Diploscapter* chromosomes are collinear

The genome sizes and predicted gene counts of both *Diploscapter* species (**Supp. Table 1**) indicated that their 2*n* = 2 chromosomes were derived from past chromosome fusion events rather than an evolutionary loss or diminution of ancestral chromosomes^8,12^. However, since the older assemblies were fragmented, we could not determine if the homologous chromosomes we see today were colinear or arose independently, which would likely give rise to very different genome architecture. Large-scale inversions and rearrangements are incompatible with meiotic recombination in nematodes^20–22^ and with few exceptions^23^, the reductional division of meiosis I depends almost universally on successful recombination.

We and others previously observed that in oocytes of *D. pachys* and *D. coronatus,* the condensed chromosomes align but do not appear to be synapsed^8,12^, raising the possibility that the loss of meiosis I in this clade is causally linked to incompatible structural rearrangements between homologous chromosomes. Instead, pairwise alignments of the four newly assembled *D. pachys* and *D. coronatus* chromosomes using nucmer^24^ revealed nearly complete collinearity with the exception of a ∼1-Mb inversion on *D. pachys* chromosome A (**Fig. 3**). This level of conserved synteny suggests that all four *Diploscapter* chromosomes derive from a single, common ancestor that arose before the divergence of these two species, and the alterations to *Diploscapter* meiosis I may be unrelated to chromosome collinearity.

**Fig. 3:**
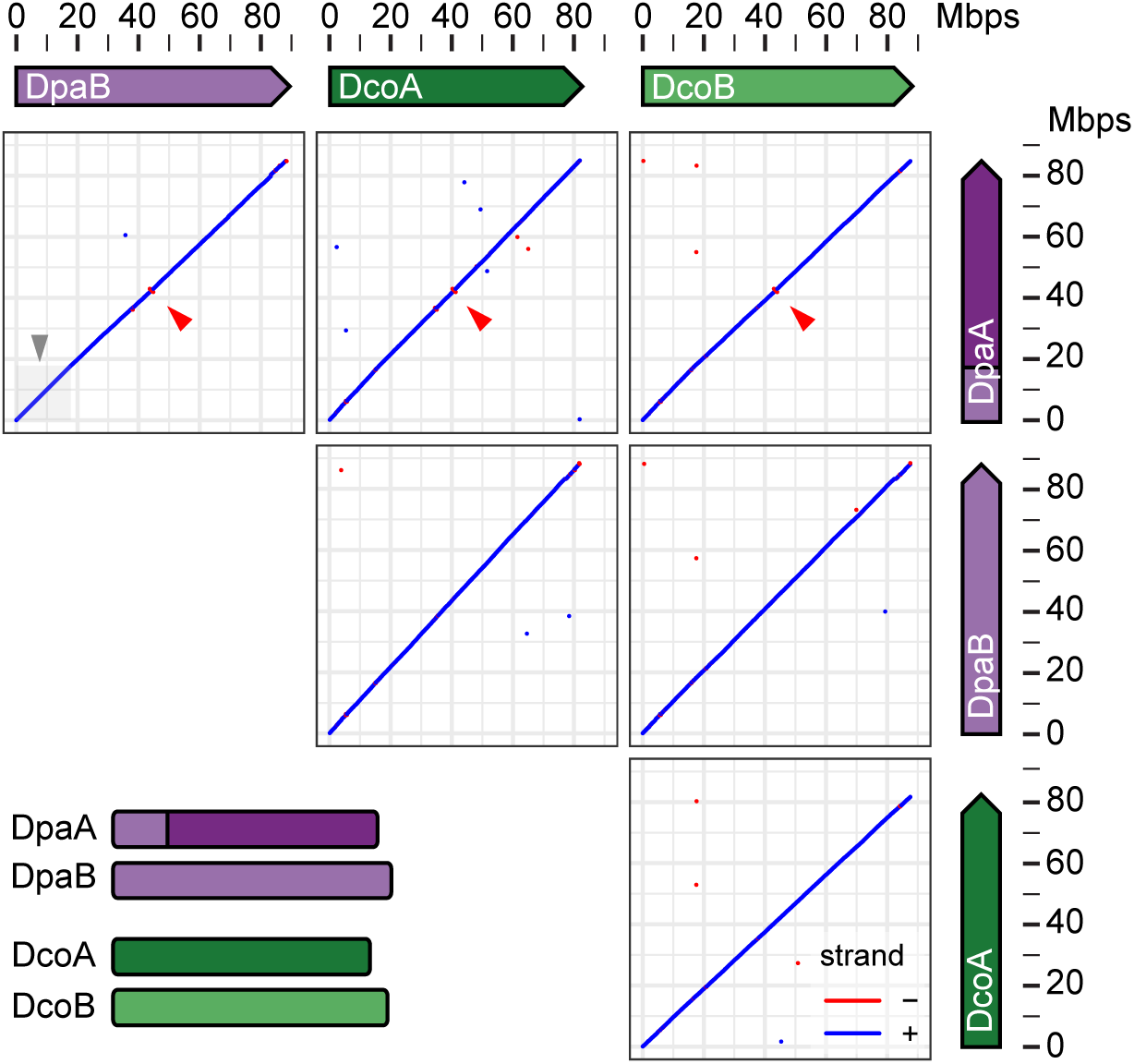
The *Diploscapter* chromosomes are colinear across their lengths. Dot plots of nucmer sequence alignments between pairs of *Diploscapter* chromosomes. Grey box in the DpaA and DpaB alignment indicates the extent of the homozygous region. Red arrowheads mark the location of a ∼1-Mb inversion unique to DpaA.

### Analysis of ancestral linkage groups reveals large-scale rearrangements from different times in the evolution of the *Diploscapter* chromosomes

Previous comparative analyses have concluded that the ancestral haploid chromosome number in rhabditid nematodes was most likely 7^13^. We therefore aimed to determine how these ancestral chromosomes may have fused to form the single haploid chromosome in *Diploscapter*. In nematodes, gene order is frequently reshuffled by intrachromosomal rearrangements, whereas translocations between chromosomes are relatively rare^13^. In the previous *D. pachys* assembly, genes within the same contig were often had *C. elegans* orthologues that mapped to a single *C. elegans* chromosome. This led to the hypothesis that the *Diploscapter* chromosome arose through a complete, end-to-end fusion of ancestral chromosomes, largely preserving their internal structure^8^.

We re-examined this hypothesis by assessing the extent of macrosynteny preserved from rhabditid ancestors to *Diploscapter*. Instead of comparing *Diploscapter* genes directly with *C. elegans*, we analysed the arrangement of seven ancestral rhabditid linkage groups known as Nigon elements^13^ that are defined by deeply conserved eukaryotic genes (BUSCO orthologs^25,26^). If the *Diploscapter* chromosomes descended from end-to-end fusion events of rhabditid ancestral chromosomes, we would expect seven clearly delineated regions, each corresponding to one predominant Nigon element. Instead, we found more than seven regions with varying levels of intermixing (**Fig. 4a**). Since rearrangements erode synteny over time, the heterogenous intermixing of Nigon elements suggests multiple, temporally distinct rearrangement events in the *Diploscapter* lineage.

**Fig. 4:**
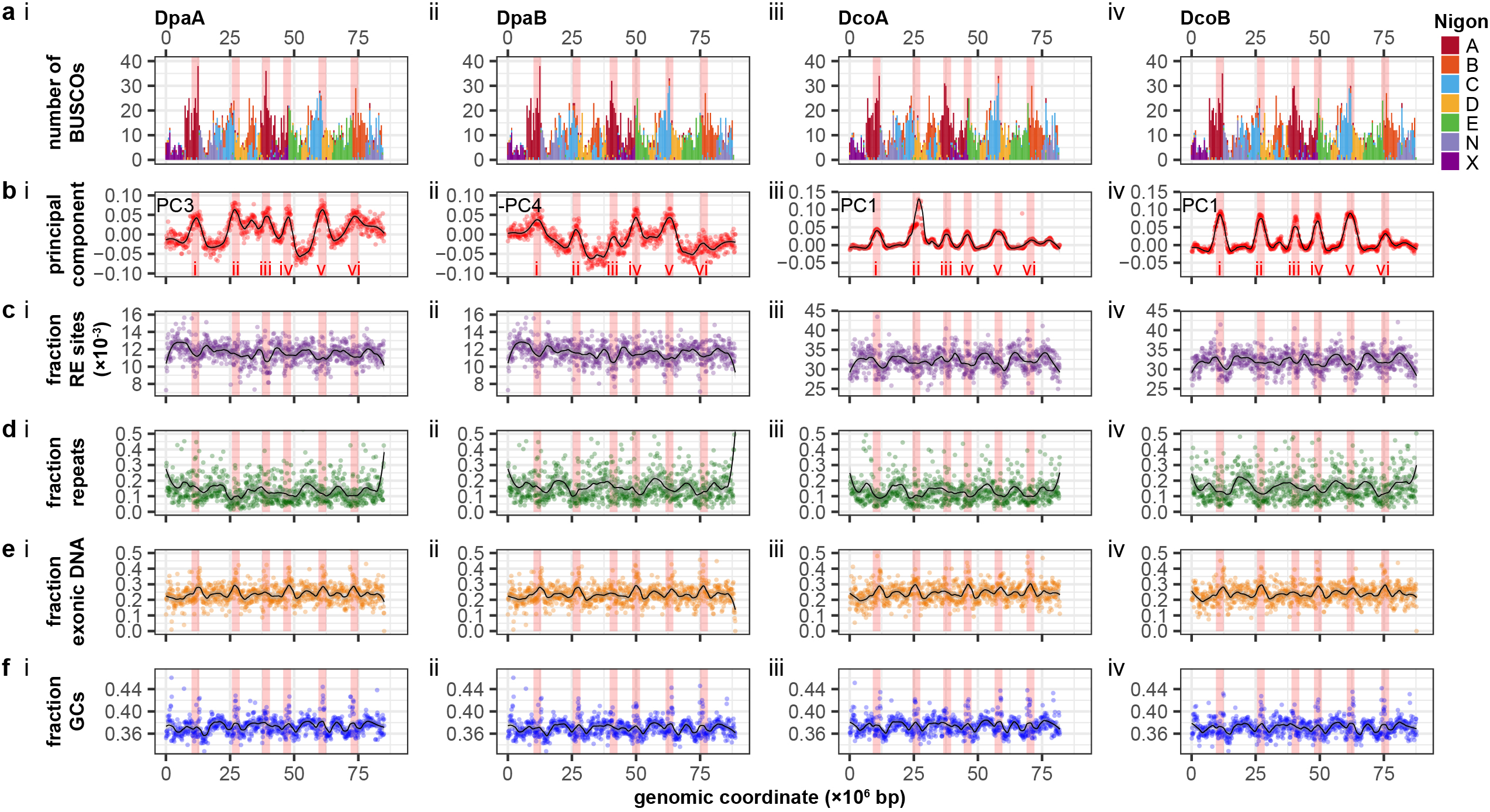
Signatures of ancestral chromosome arrangements. **a-f,** Various chromosome traits along DpaA (i), DpaB (ii), DcoA (iii) and DcoB (iv) chromosomes. The six high frequency contact regions are highlighted in red. **a**, Arrangements of ancestral linkage groups (Nigon elements) with 500,000-bp bins. **b**, High frequency contact (HFCs) regions revealed by principal component analysis on PoreC (*D. pachys*) or HiC (*D. coronatus*) data. c, Density of restriction enzyme sites for PoreC (DpaA, DpaB) and Arima Hi-C (DcoA, DcoB) in 100,000-bp bins. **d**, Density of repetitive DNA in 100,000-bp bins. **e**, Density of exonic sequences in 100,000-bp bins. **f**, GC content in 100,000-bp bins.

### *Diploscapter* chromosomes retain vestiges of ancestral chromosome features

To detect signatures of the ancestral chromosomes immediately before the karyotype reduction to a single pair of chromosomes (2*n* = 2), we scanned for molecular traits consistent with historical fusions (**Fig. 4**). For comparison, we examined genomic features in a recently published assembly of a more distant dioecious relative with 6 chromosomes (2n = 12), *Pristionchus exspectatus*^27^ (**Supp. fig. 2**). In addition to largely preserving the macrosynteny of Nigon elements (**Supp. fig. 2a**), rhabditid chromosomes typically exhibit distinct functional differences between arms and more central regions^28^, which have lower repeat and GC content, and higher exonic DNA (**Supp. fig. 2b–d**, red highlights). Moreover, frequently interact with each other in the nucleus^27,29^ (**Supp. fig. 2e**, red highlight), reminiscent of the ‘A’ compartments observed in chromosome contact maps of mammalian cells^30^. Remarkably, hallmarks of chromosome centrality can persist after ancestral linkage groups fuse: the *Pristionchus exspectatus* X chromosome – a recent fusion of Nigon N and X – contains two regions with these characteristics^27^ (**Supp. fig. 2**).

Remarkably, chromosome contact maps of both *D. pachys* and *D. coronatus* reveal at least six long-range high-frequency contact (HFC) areas reminiscent of ancestral chromosome central regions (**Fig. 1b**, red arrowheads), which are also supported by principal components analysis (**Fig. 4f**). The number of HFCs is consistent with the most common karyotypes observed in Rhabditida (5 ≤ *n* ≤ 7) and is not an artefact of restriction site density (**Fig. 4c, Supp. fig. 3a**). Five of the six HFCs (i, ii, iv, v and vi) share characteristics with chromosome centres of other nematode species, tending toward lower repetitive DNA and higher exonic DNA content (**Fig. 4d-e, Supp. fig. 3b-c**). Intriguingly, these five HFCs also coincide with Nigon domain boundaries. However, unlike chromosome centres in *Pristionchus expectatus* (**Supp. fig. 2d**), these *Diploscapter* HFCs are not more AT-rich than other regions (**Fig. 4f, Supp. fig. 3d**). Altogether, the frequent long-range contacts, lower repeat content, and higher exonic DNA content point to at least six centre-like regions along the *Diploscapter* chromosome.

### *Diploscapter* chromosomes show evidence of sex and introgression after the speciation of *D. pachys* and *D. coronatus*

The degree of nucleotide identity between *Diploscapter* chromosomes should reflect their relatedness and reveal their evolutionary history. Excluding the homozygous region of the *D. pachys* genome, where homologs A and B share 100% sequence identity, the heterozygous region of *D. pachys* homolog A (DpaA) and the corresponding region of *D. coronatus* homolog A (DcoA) are more similar to each other (∼97% identity) than all other within– and between-species pairwise comparisons (∼93% identity) (**Fig. 5a**). To understand the evolutionary history of these chromosomes, we used the conserved nematode gene orthologs (BUSCOs^25,26^) and concatenated their coding sequence alignments to perform a maximum likelihood analysis^31,32^ of their evolutionary relationships.

**Fig. 5:**
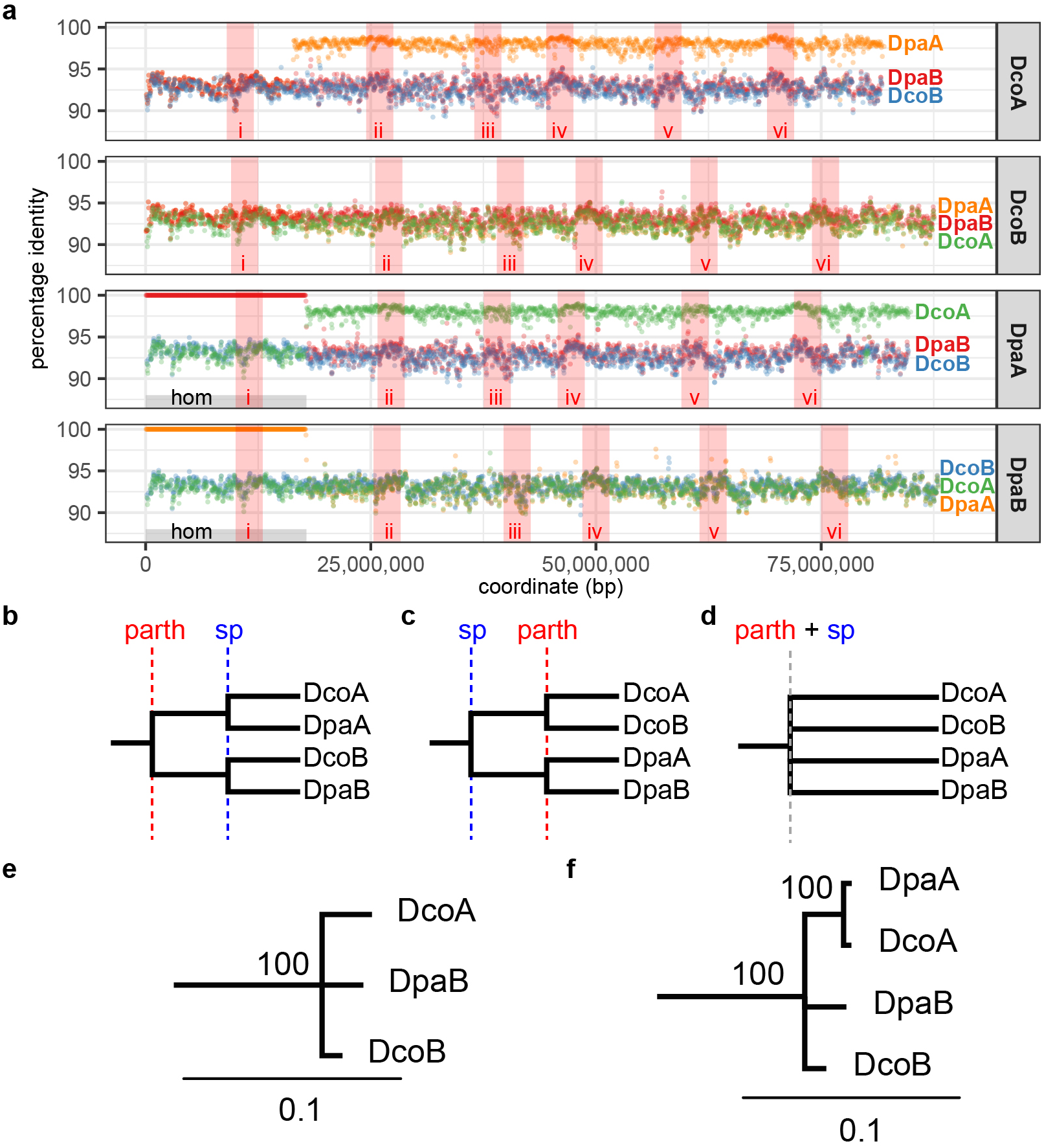
Patterns of sequence identity suggest *D. pachys* and *D. coronatus* A chromosomes are more closely related. **a**, Percentage sequence identity when a *Diploscapter* chromosome (label on the right) is compared to the other chromosome (colored dots on plot). HFCs are shown as red vertical bars. The *D. pachys* homozygous region is at the left end (grey). **b**, Pattern of divergence if the emergence of parthenogenesis preceded the speciation of *D. coronatus* and *D. pachys*. **c,** Pattern of divergence if speciation preceded the emergence of parthenogenesis. **d**, Pattern of divergence if the emergence of parthenogenesis coincided with the speciation of *D. coronatus* and *D. pachys*. **e**, The observed pattern of divergence of the *Diploscapter* chromosomes inferred from RAxML analysis using orthologous genes in in the *D. pachys* homozygous region. The divergence most resembles **d**. **f**, The observed pattern of divergence of the *Diploscapter* chromosomes inferred from RAxML analysis using orthologous genes in in the *D. pachys* heterozygous region. The A homologs are more closely related to each other.

We considered three scenarios for how the sequences of the four *Diploscapter* chromosomes could have diverged, depending on when parthenogenesis – which we equate here with loss of recombination – arose in relation to the speciation event (**Fig. 5b-d**). If parthenogenesis emerged prior to speciation (**Fig. 5b**), and the two chromosomes in the stem species began diverging due to lack of recombination, then we may expect the orthologous chromosomes (e.g. DpaA and DcoA) to share greater sequence similarity than the homologous chromosomes (e.g. DpaA and DpaB). If speciation preceded parthenogenesis (**Fig. 5c**), then genetic exchange between homologs would presumably have been possible until the (independent) onset of parthenogenesis in each lineage, giving rise to greater within-than between-species similarity. If instead these events coincided (**Fig. 5d**), all *Diploscapter* chromosomes should be equally divergent, assuming similar nucleotide substitution rates in the two lineages.

Our analysis reveals a pattern of sequence divergence that is incompatible with any of these three models. Consistent with their higher sequence identity (**Fig. 5a**), a pairwise comparison of the two *Diploscapter* A chromosomes shows fewer substitutions per site than any other pair of *Diploscapter* chromosomes, which all show a similar rate of substitutions (**Fig. 5e**). Thus, the common ancestor of the A chromosomes is more recent than that of all four chromosomes. Finally, in *D. pachys* only, a single homogenizing event led to the replacement of 17.8 Mb of chromosome A with material from chromosome B, giving rise to the homozygous region in the *D. pachys* genome.

## Discussion

The new genome assemblies of *D. pachys* and *C. coronatus* described here – based on a combination of long-read sequencing and chromosome contact maps – enable us to refine our previous hypotheses regarding their genome architecture and its relationship to the evolution of parthenogenesis in this clade. By examining the distribution of ancestral linkage groups, or Nigon units, along the *Diploscapter* chromosomes, we discovered that while many regions remain strongly enriched for a single Nigon unit, they have undergone extensive reorganization (**Fig. 4a**). Notably, the approximate boundaries of contiguous domains with a dominant Nigon signature often coincide with high-frequency contact regions (HFCs) in chromosome contact maps (**Fig. 4a**). The ancestral linkage groups can be reconstituted *in silico* by rearranging these domains to produce an adjacency map of Nigon segments in which HFCs are centrally located within each linkage group (**Fig. 6a.i**). Based on the observed level of intermixing – and presuming that intrachromosomal rearrangements occur at higher frequency – this linkage arrangement suggests a likely intermediate karyotype (2*n* = 8) in the evolution of the *Diploscapter* lineage, in which the A-X and B-C-N telomere-telomere fusions would have occurred at an earlier time, and further fusion events would then have reduced the chromosome count to the current 2*n* = 2 karyotype (**Fig. 6a.ii-iv**). A similar proposal involving A-X and B-C fusions has been explored previously^8^.

**Fig. 6:**
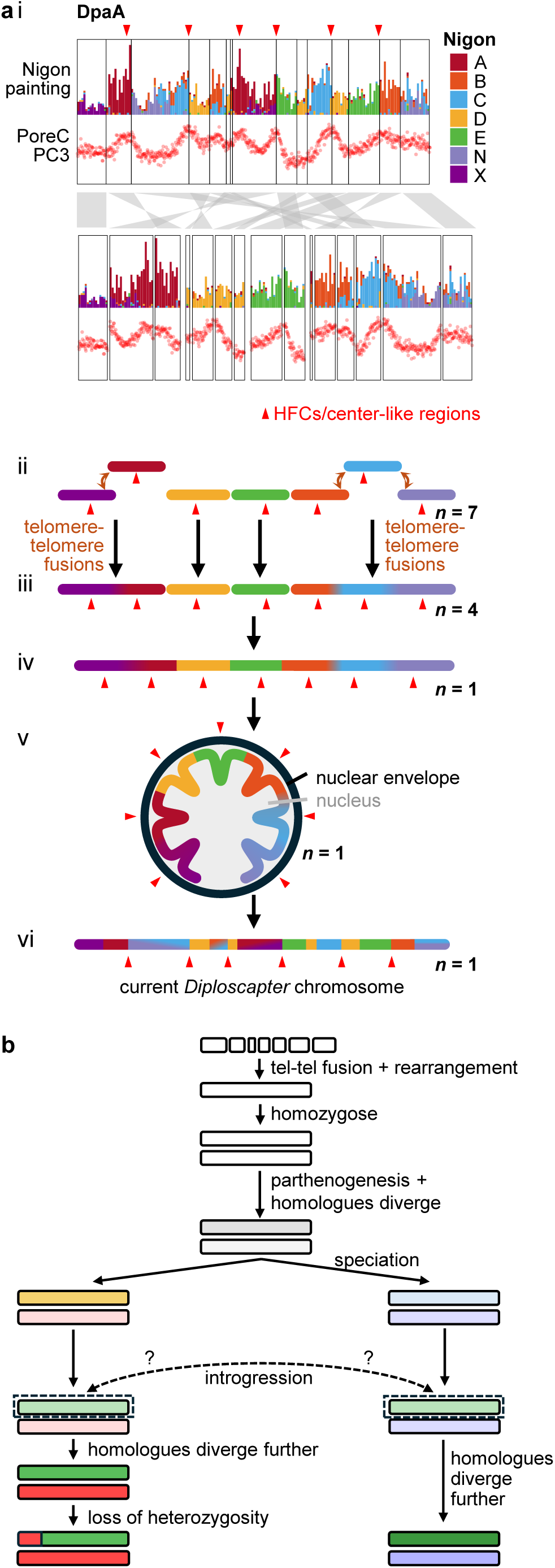
Models for the evolution of the *Diploscapter* chromosomes. **a**, A model for the formation of the *Diploscapter* fusion chromosome. **i,** Rearranging contiguous Nigon blocks on DpaA reveals a linkage arrangement which may have existed during the evolution of this species. When the principal component values from PoreC are rearranged in the same way, regions with high contact (red arrowheads) tend to line up at the centre of each rearranged Nigon block. **ii,** The ancestral Rhabditid chromosomes corresponding to the Nigon elements. The A-X and the B-C-N end-to-end fusions occurred early in the evolution of *Diploscapter*. **iii**, Local rearrangements along the fusion chromosomes partly intermix the segments. This karyotypic arrangement is reflected in panel i (bottom). **iv**, Further telomere-telomere fusions occurred to reduce the karyotype to 2*n* = 2. **v**, One last round of rearrangements might have involved regions of close contact: centre-like regions exchanged with each other, and arm-like regions exchanged with each other. **vi**, the current *Diploscapter* chromosome, which took shape before the speciation of *D. pachys* and *D. coronatus*. **b,** Possible evolutionary paths of *D. pachys* and *D. coronatus*, whose more similar A homologs may be due to a rare mating event. In this scenario, ancestors of *D. pachys* and *D. coronatus* embarked on their separate paths after the appearance of parthenogenesis, but a rare mating between the two replaced the native A homolog.

The new data from the HiC and PoreC contact maps generated for the new assemblies allow us to posit that historical chromosome centres maintained frequent contacts even after the formation of the single large chromosome (**Fig. 6a.v**). This proximity may have permitted additional rearrangements among them that could explain the observed congruence of HFCs with the current Nigon domain boundaries (**Fig. 6a.vi**). The evolution of *Diploscapter* chromosomes by structural rearrangements is therefore likely constrained by the topology of chromatin within the nucleus. Similarly, somatic structural variant sites in human cancer cell lines often coincide with points of interaction between topologically associated regions^33^. Thus, rearrangements between discontinuous euchromatic regions that are constrained by their physical proximity in the nucleoplasm may be a general property of eukaryotic chromosome evolution. It will be informative to generate genome assemblies for additional species in the *Protohabditis* group whose karyotypes may represent frozen intermediates in the fusion process.

The chromosome fusions are almost certainly linked to the short telomeric repeats and long subtelomeric arrays revealed by our current study, though we do not yet know the precise sequence of events or causal relationships that led to the current karyotype. Telomeres are susceptible to shortening because the DNA replication machinery cannot synthesize the extreme 5’ ends of double-stranded DNA^34,35^.

Eukaryotic telomeres are usually bound by proteins that control access to the DNA, preventing DNA repair proteins from erroneously joining chromosome ends^36^ or recruitment of telomerase to lengthen the telomeres^37^. Both *Diploscapter* species appear to be missing one such telomere-binding ortholog with an OB-fold^8^. Moreover, the *Diploscapter* ortholog of another telomere-binding protein, TEBP-1/2, lacks the PBR domain^38,39^ that interacts with telomeric OB-fold proteins (**Supp. fig. 4**). Thus, the ancestral mechanism for maintaining telomere length involving an OB fold protein has been altered or lost in *Diploscapter*.

Other than telomerase, one mechanism to counteract telomere shortening is the Alternative Lengthening of Telomeres (ALT), which uses proteins associated with homologous recombination and break-induced repair to copy new sequences to the chromosome ends^40^. Sometimes this results in repetitive motifs different from those maintained by telomerase. Curiously, the structure of *Diploscapter* subtelomeric repeats (**Fig. 2, 3d**) resembles chromosome ends shaped by ALT in at least four instances: *C. elegans* telomerase-deficient mutants^41,42^, wild *C. elegans* isolates^43^, wild *C. briggsae* isolates^44^, and a line of telomerase-RNA-deficient mouse embryonic stem cells^45,46^. The *Diploscapter* subtelomeres, therefore, carry sequence signatures similar to a specific type of templated ALT (“TALT”) event^38^.

We hypothesize that the evolutionary loss of the telomeric OB-fold protein prevented the efficient recruitment of telomerase to chromosome ends, resulting in short telomeric repeat arrays that were occasionally lost (**Supp. fig. 5a-b**). At different times during the evolution of this lineage, such losses would have given rise to end-to-end fusions, leading to a step-wise reduction in chromosome number that ultimately resulted in a single pair of 80-90 Mb chromosomes (**Fig. 6a.ii-iv**, **Supp. fig. 5c-d**). Fusion products harbouring additional copies of ancestral chromosomal units would be unlikely to survive due to gene dosage constraints (**Supp. fig. 5e**). Without robust maintenance of telomeric repeats, an alternative mechanism would be needed to prevent the further degradation of chromosome ends. We propose that TALT could maintain the ∼3 kb subtelomeric repeats, potentially along with adjoining telomeric repeats (**Supp. fig. 5f**). In the absence of recombination, sequences at the different chromosome ends would diverge over time (**Supp. fig. 5g**), while recurrent activation of ALT would replenish the subtelomeric repeat arrays, resulting in the long *Diploscapter* subtelomeres we see today (**Supp. fig. 5h**).

Chromosome evolution driven by long-term telomere dysfunction is not unique to *Diploscapter*. In Dipteran species, the loss of the telomerase gene and the use of transposons for telomere homeostasis are hypothesised to drive the rapid sequence changes to telomere-binding proteins^47,48^, analogous to sequence changes in the *Diploscapter* TEBP orthologue (**Supp. fig. 4)**. *Strongyloides* nematodes have also lost the telomerase gene^49^, and similar to the two *Diploscapter* species described here, several *Strongyloides* species have a reduced chromosome count resulting from fusion events, a history of rearrangement among the ancestral linkage groups, and chromosome ends with long non-canonical repeats likely expanded through ALT^49,50^. These convergent traits demonstrate that long-term telomere dysfunction can be a potent driver for chromosome evolution of holocentric nematode chromosomes^28^. This is particularly salient because the classic Breakage-Fusion-Bridge (BFB) cycle^51^ – a major cause of large-scale rearrangements and genomic instability in cancer cells and, more generally, animals with point centromeres – does not affect the mitotic success of holocentric chromosomes.

Within the *Diploscapter/Protorhabditis* clade, several other parthenogenetic nematode species have been identified^8,52–54^, but their phylogenetic relationships have not been clearly resolved. Our new genome assemblies allowed us to consider the relative timing of speciation and the emergence of parthenogenesis in *D. pachys* and *D. coronatus*. Sequence comparisons revealed that the A homologs in these two species must have diverged at a later time than the point at which their shared ancestral A chromosome diverged from the *D. pachys* and *D. coronatus* B homologs (**Fig. 5f**). The most parsimonious explanation for this is that the onset of parthenogenesis roughly coincided with the speciation event separating *D. pachys* and *D. coronatus*, but a later introgression event brought the same ancestral A homolog into both linages (**Fig. 6b**). How this genetic exchange occurred is not known, given that few or no *Diplosocapter* males have been observed in the wild or in the laboratory, and *Diploscapter* species readily reproduce by ameiotic parthenogenesis^53–61^.

Generating end-to-end genome assemblies for several other parthenogenetic *Diploscapter* and *Protorhabditis* isolates would enable a more comprehensive analysis of genetic exchanges between different isolates and among different chromosomes. Determining the frequency and the timeline of these exchanges would provide new insights into the mechanisms by which they may have occurred. Perhaps rare genetic exchanges provide a mechanism for *Diploscapter* parthenogens to limit the accumulation of excessive deleterious mutations and thereby escape Muller’s rachet? Additionally, *Diploscapter* may be a species complex where many diverse but related species occasionally interbreed and produce sub-fertile lineages whose genetic innovations help them circumvent the requirement for males in reproduction.

These outstanding questions underscore the importance of broader genomic sampling to investigate and resolve their evolutionary history, which could lead to more general insights into genome evolution and a deeper mechanistic understanding of reproductive plasticity.

## METHODS

### Strain maintenance

*Diploscapter pachys* PF1309 animals were grown on a cholesterol-enriched medium with high agarose content to reduce burrowing (51.3 mM NaCl, 1 mM CaCl_2_, 1 mM MgSO_4_, 25 mM KPO_4_ pH 6, 100 µg/mL cholesterol, 3% w/v agarose) and fed on a diet of *E. coli* OP50-1 enriched with cholesterol (grown in LB broth with saturated cholesterol), as *D. pachys* animals appear to thrive better on a higher dietary cholesterol than other nematodes (unpublished observations). For *Diploscapter coronatus* strain PDL0010, we cultured the animals at 20°C on nematode growth media (NGM) plates seeded with *E. coli* HB101. A layer of water on the plate reduced burrowing.

### Harvesting nematodes for DNA extraction

We harvested the nematodes by washing the plates several times with cold M9 into 50 ml Falcon tubes, which we then centrifuged at 4000 rcf for 8 min. We discarded the supernatant and washed the nematodes a further two times using M9 supplemented with 0.01% Tween. We then performed two sucrose flotation wash steps to remove bacteria and other contamination. For the sucrose wash, we transferred the worms in a 15 ml Falcon tube and resuspended the worm pellet in 5 ml cold M9. We added 5 ml of 60% chilled sucrose solution and centrifuged the suspension for 5 min with 3400 rcf at 4°C. We collected the nematode from the top layer and washed with 14 ml M9 supplemented with 0.01% Tween. After the two sucrose washes, we performed a final wash using PBS buffer. We divided the worms into 1.5 ml DNA LoBind® Tubes (Eppendorf) before flash freezing in liquid nitrogen and storing at –70°C.

### Sequencing *Diploscapter* material

*D. pachys* PF1309 DNA was extracted using the MagAttract HMW DNA kit from Qiagen per the manufacturer’s instructions. Genomic DNA library was constructed using the DNA ligation kit from Oxford Nanopore Technologies (Oxford, UK) following the manufacturer’s instructions. The library was sequenced on a Oxford Nanopore R9.4.1 flowcell (FLO-MIN106) and basecalled using Guppy (v 4.0.11). Subsequently, library adapters were trimmed using Porechop^62^ (v.0.2.4) and reads were filtered by length (8000 bps, –l 8000) and by quality (8 or better, –q 8) using NanoFilt^63^ (v.2.6.0).

For *D. coronatus* PDL0010, we performed two independent high molecular weight DNA extractions: (1) Employing the Nanobind Tissue Big DNA Kit and the Buffer NL (Circulomics) on a 100mg worm pellet using the “Nanobind High Molecular Weight C. elegans DNA Extraction Protocol” with the following modifications: we added 20 µl Proteinase K and 150 µl Buffer NL to the frozen pellet and disrupted the tissue with the BioMasher II. We eluted the DNA from the nanobind disk overnight in 200 µl EB buffer. We then mixed the extracted DNA several times with a bore pipette tip before proceeding to QC measurements. We sheared the DNA to an average size of 14 kb with a Megaruptor 3 (setting 29) (Diagenode). The sheared DNA was SPRI cleaned with 1.8x of AMPure XP beads (Beckman Coulter). (2) Using the MagAttract HMW DNA kit (Qiagen) with the following modifications: We used a pellet of nematodes of 80 mg. The lysis mix was prepared and placed on ice: 200 µl PBS, 20 µl ProteinaseK (Qiagen), 4 µl RNase A (Qiagen), 150 µl AL buffer (Qiagen). We added 75 µl of the lysis buffer mix to the frozen nematode pellet and used a BioMasher II to disrupt the pellet. We added the remaining lysis buffer and mixed with a wide bore tip. We transferred the lysis solution to a 2 ml DNA LoBind® Tube (Eppendorf) and digested overnight at 45°C mixing at 600 rpm in a ThermoMixer C (Eppendorf). We added 15 µl of MagAttract Suspension G before adding 280 µl Buffer MB. We eluted the MagAttract beads twice using 200 µl of Buffer AE in each elution step. We incubated the second elution mix at 25°C with 1000 rpm for 3 min in the ThermoMixer C before transferring the elution liquid to a new 1.5 ml LoBind microtube. We sheared the DNA to an average size of 10.3 kb with a Megaruptor 3 (setting 29) (Diagenode). The sheared DNA was SPRI cleaned with 1.8x of AMPure XP beads (Beckman Coulter).

Two *D. coronatus* PacBio libraries were prepared by the Scientific Operations: Sequencing Operations core at the Wellcome Sanger Institute using the PacBio Low DNA Input Library Preparation Using SMRTbell Express Template Prep Kit 2.0. Both libraries were sequenced on a single PacBio Sequel IIe flow cell.

### Sequencing of proximity-ligated DNA

A sequencing library for *D. pachys* Pore-C was generated by extracting DNA crosslinked to chromatin that was subsequently digested by a restriction enzyme and religated. Briefly, *D. pachys* animals suspended in M9 buffer (22.0 mM KH_2_PO_4_, 42.3 mM Na_2_HPO_4_, 85.6 mM NaCl, 1 mM MgSO_4_) were flash-frozen in liquid nitrogen and ground up using mortar and pestle to release cellular contents and nuclei. Chromatin and cellular components were crosslinked using 1% w/v formaldehyde in PBS (137 mM NaCl, 2.7 mM KCl, 8 mM Na_2_HPO_4_, 2 mM KH_2_PO_4_, pH 7.4) at room temperature for 10 minutes. After quenching the formaldehyde with 1% w/v glycine and the removal of formaldehyde, nuclei were permeablised using 0.2% v/v IGEPAL (with 10 mM Tris-HCl and 10 mM NaCl, pH 8) supplemented with protease inhibitor cocktail P8340 (Sigma) and heat-treated at 65 °C for 1 hr. Chromatin was denatured using 0.1% w/v sodium dodecyl sulfate (SDS) and the DNA was then digested by *Nla*III (New England Biolabs) at 37°C overnight. *Nla*III was subsequently deactivated by heat (65°C for 20 minutes) and *Nla*III-digested ends were allowed to anneal by slowly cooling to room temperature before ligation using T4 DNA ligase (New England Biolabs). Finally, proteins were digested from the crosslinking ligation reaction by incubating with 1 μg/μL proteinase K (Invitrogen, Waltham, MA) at 56°C for 18 hours. DNA from this preparation was purified using standard phenol-chloroform extraction and resolubilized with TE buffer. The purified, proximity-ligated DNA was ligated to Nanopore adapters per the manufacturer’s instructions. The library was then sequenced on an Oxford Nanopore R9.4.1 flowcell (FLO-MIN106) and basecalled using Guppy (v 6.0.7).

*D. coronatus* Hi-C library preparation and sequencing were performed by the Scientific Operations: Sequencing Operations core at the Wellcome Sanger Institute. A 25 mg pellet of mixed-stage nematodes was processed using the Arima Hi-C version 2 kit following the manufacturer’s instructions. An Illumina library was prepared using the NEBNext Ultra II DNA Library Prep Kit and sequenced on one-eighth of a NovaSeq S4 lane using paired-end 150 bp sequencing.

For both species, visualisation of the HiC and PoreC data was done in Juicebox (v1.11.08)^64^, or the HiCExplorer suite (v3.7.2)^65^. Tools in the HiCExplorer suite was used for principal component analysis of the HiC and PoreC matrices.

### Assembly and scaffolding of *Diploscapter* genomes

A draft of the *D. pachys* genome was assembled *de novo* using Nanofilt-filtered reads and Canu^66^ (v.2.1) with “heterozygous” settings (“batOptions=-dg 3 –db 3 –dr 1 –ca 500 –cp 50”) recommended by the authors of Canu. Canu was chosen as it assembled the two haplotypes separately, while assemblers such as flye^67^ and wtdbg2^68^ collapsed the two haplotypes (data not shown). Contigs derived from bacterial contamination were identified by BLASTN^69^ against the RefSeq database of all known bacterial genomes and removed, leaving 53 contigs. One round of contig correction was done using Pilon^70^ (v 1.24) and the published *D. pachys* Illumina genomic reads^8^ mapped to the 53 contigs (SRA accession SRX5194681-SRX5194684). One of these contigs was the mtDNA, which was 13393 bps after Pilon correction and manual curation. Additional rounds of correction by Pilon did not noticeably improve the assembly (data not shown).

*D. pachys* Pore-C sequencing data were processed using the Pore-C Snakemake pipeline (https://github.com/nanoporetech/Pore-C-Snakemake), which mapped them onto the *D. pachys* contigs assembled by Canu and corrected by Pilon. Scaffolding of the contigs was done using YaHS^71^ (v1.1) and the contact matrices from the Pore-C Snakemake pipeline. After scaffolding, significant sequence overlaps between adjacently positioned contigs were checked manually and trimmed. The result was a scaffolded genome assembly with 11 non-mitochondrial scaffolds: 2 larger than 70 Mbps, which represented the nuclear chromosomes; and 9 smaller contigs amounting to about 500,000 bps – less than 0.3% of the total sequence – which could not be placed on the nuclear chromosome scaffolds and were thus left out of the final scaffolded assembly.

The span of the D. pachys homozygous region was determined by mapping Nanofilt-filtered genomic Nanopore reads back to the scaffolded assembly. The homozygous region was initially only assembled once by the Canu assembler and scaffolded to the DpaA chromosome but not the DpaB chromosome. On this DpaA scaffold, the heterozygous-homozygous boundary was readily apparent at nucleotide position 17,839,851, where the Nanopore read mismatches with a frequency ≈ 0.5 abruptly appear to the left of this position, and abruptly disappear to the right of this position. Thus, the terminal 17,839,851 bps of this scaffold was added to the corresponding heterozygous-homozygous boundary on DpaB, yielding the final curated DpaA (85,103,150 bps) and DpaB (88,703,182 bps) chromosomal scaffolds whose first 17,839,851 bps are identical. For certain analyses where the homozygous region would be redundant (such as gene annotation), the 17,839,851-bp homozygous region was left out of DpaB scaffold.

To assemble the *D. coronatus* genome, we removed adapter sequences from the PacBio HiFi data using HiFiAdapterFilt^72^. We used Jellyfish 2.3.0^73^ to count k-mers of length 31 in each read set and GenomeScope 2.0^74^ to estimate genome size and heterozygosity. We first generated a primary and alternate assembly from the PacBio HiFi data using hifiasm v0.16.1-r375^75^. We randomly subsampled 10% of the Hi-C reads using samtools^76^ v1.14 and aligned them to the hifiasm primary assembly using bwa-mem 0.7.17-r1188^77^, filtered out PCR duplicates using picard 2.27.1-0 (available at http://broadinstitute.github.io/picard/), and scaffolded the assembly using YaHS^71^. We ran BlobToolKit 2.6.5^78^ on the scaffolded assembly and used the interactive web viewer to manually screen for scaffolds derived from non-target organisms. We removed 1,032 contigs corresponding to various Proteobacteria, Bacteroidetes, and Actinobacteria species. After removing contaminants, we used MitoHiFi 2.2^79^ to extract and annotate the mitochondrial genome. We removed residual haplotypic duplication from the assembly using purge_dups 1.2.5^80^ and scaffolded the purged primary assembly using Hi-C data, as previously described. We used seqkit^81^ and BUSCO 5.2.2^25,26^ with the “nematoda_odb10” lineage to calculate assembly metrics and assess biological completeness. We assessed base-level accuracy and k-mer completeness using Merqury 1.3^82^.

The final *D. coronatus* assembly was checked for residual contamination and corrected using BlobToolKit^78^, and FCS^83^. We removed four scaffolds that corresponded to protists, bacteria, and fungi. Manual curation was performed using JBrowse 2^84^, HiGlass^85^ and PretextView (available at https://github.com/wtsi-hpag/PretextView), where we made eight breaks, four joins, and removed four haplotigs.

### Chromosome-wide sequence comparisons using nucmer

Nucmer (v 4.0.0rc1), a part of the Mummer4 suite of programs^24^, was used to align two chromosome scaffolds against each other for all pairwise comparisons, each generating an intermediate ‘delta’ file which recorded all local alignments between the pair of chromosomes. The program delta-filter, also part of the Mummer4 suite, was used to eliminate short, spurious, and repetitive local alignments (options –qr – l 10000) in these delta files. This generated a list of the longest consistent sets of local alignments longer than 10,000 bps. Custom scripts in Python and R were used to parse through the alignments and plot sequence identities as a dot plot.

To determine the percent sequence identity between two chromosomes being aligned, we first manually reverted the ∼1 Mbp inversion on DpaA to its likely uninverted ancestral state (coordinates 41,921,280 to 42,998,403), then re-ran nucmer as above in all pairwise combinations. Delta-filter was then run in ‘global’ mode (option –g) on each pairwise delta file, retaining only the longest mutually consistent set of non-inverted and non-repetitive local alignments. The Mummer4 utility show-coords (options –rT) was used to list the start and end coordinates of each alignment, while the Mummer4 utility show-snps (options –CrT) was used to list the alignment mismatches, which can be single-nucleotide substitutions or small indels.

We wrote a custom Python script to run show-coords and show-snps and use their outputs to calculate the percentage sequence identities across the entire chromosomes in genomic windows of 100,000 bps. The sequence identity is defined as [number of nucleotide mismatches within the alignments in the window]/[the span of all local alignments in the window while accounting for indels] * 100%. In our calculations, we did not include any nucleotides that nucmer could not align. Because not every nucleotide was counted in each window, we chose to omit the 100,000-bp windows where fewer than 70,000 bps (70%) were aligned by nucmer. Finally, a custom R script was used to plot the sequence identities calculated at all windows.

### Genome annotation and BUSCO search for conserved eukaryotic orthologs

For both species, ribosomal RNA genes were predicted using RNAmmer^86^ (v1.2). Protein-coding genes were predicted by the Braker2 pipeline^87^ using Hisat2^88^ mapping the published RNAseq reads^8,12^ (SRA run accessions: *D. pachys* SRX5194679 and SRX5194680, *D. coronatus* DRX026964). For both *D. pachys* and *D. coronatus* BRAKER2 runs, we disabled *de novo* prediction of any genes without RNAseq support by setting the intron “malus” value to 0 in the Augustus configuration file rnaseq.cfg (see https://github.com/Gaius-Augustus/BRAKER/issues/667), and additionally removing any predicted genes whose FPKM value is 0 using StringTie^89^ (expression estimation mode, option –e). In *D. pachys*, stranded RNAseq libraries were used to annotate each DNA strand separately, which in some cases improved the prediction accuracy of genes located inside the introns of other genes. *D. coronatus* RNA-seq libraries were unstranded and thus this was not done. We did not mask the repetitive elements in the genome prior to gene annotation – by visual inspection, RepeatMasker^90^ masked many regions with clear RNA-seq support.

We performed BUSCO searches^25,26^ (v 5.4.2) for conserved gene orthologs to estimate the completeness of the assemblies as well as to visualize ancestral linkage groups (Nigon painting^13^). For each *Diploscapter* chromosome, the BUSCO searches attempted to find a set of 3131 proteins defined for the Nematoda lineage (-l nematoda_odb10) in the sets of proteins (-m protein) predicted by Braker2 above. Proteins were matched with the corresponding encoding loci using the annotation (gtf/gff) files generated above. Nigon painting was then performed as previously described^13^.

### Determining the GC content, the exonic DNA content, the repeat content, and the HiC/PoreC principal components

Calculating GC content, exonic DNA content and repeat content were done using custom Python scripts, in non-overlapping 100,000-bp windows. Repeat content per 100,000 bps was estimated using the RepeatScout^91^ (v 1.0.6) called by a custom Python script, based on a protocol described by Yoshida & al^27^. Each individual homologous chromosome was analyzed separately with this protocol. Briefly, the script first analyzed a haploid genome by running RepeatScout’s build_lmer_table using an l-mer size of 14 (option –l 14). Then, the script called RepeatScout using the identical l-mer size (option –l 14) and the l-mer table from the previous step to identify repeats and to generate a repeat library. The script then ran RepeatScout’s filter-stage-1.prl script to eliminate any repeats shorter than 50 bps and with over 50% low complexity as identified by TRF^92^ (v 4.09), thus generating a curated repeat library. RepeatMasker was called using this curated repeat library and the genome file to generate a repeat annotation file (.gtf), while a separate function in the script determined the repeat occurrence to limit annotations to only those occurring more than 5 times in the genome. Using this annotation file, the script determined the repeat occupancy in every non-overlapping 100,000-bp window in a genome.

Principal component analysis (PCA) on the chromatin contacts matrix was performed using the hicPCA program from the HiCExplorer suite (v 3.7.2)^65^. The analysis of *D. pachys* chromatin contact was complicated by the presence of a homozygous region in the genome. To circumvent this problem, *D. pachys* PoreC reads were first mapped with PoreC Snakemake (https://github.com/nanoporetech/Pore-C-Snakemake) to an assembly where the homozygous region was represented only once. With the resulting .pairs file, we wrote a custom Python script to randomly assign the reads mapping to the homozygous region to either DpaA or DpaB. The new .pairs file was sorted using pairtools^93^ (v 1.0.2, https://github.com/open2c/pairtools), converted into a .hic file using juicertools (v 1.19.02) from the Juicebox suite^64^, and visualized using Juicebox or HiCExplorer^65^. The *D. pachys* PoreC matrix had much more noise, and the long-range inter– and intra-chromosomal contacts discussed in this work are seen in PC3 (DpaA) or PC4 (DpaB) instead of PC1 in DcoA/DcoB.

### Inferring maximum likelihood phylogenetic trees using concatenated BUSCO coding sequences and RAxML

We developed a Python script to automate the multiple sequence alignment and the construction of maximum likelihood trees. Treating each *Diploscapter* chromosome as a haploid genome, our script first identified a subset of BUSCO genes that can be found on every sequenced *Diploscapter* chromosome and in the genomes of *Caenorhabditis elegans* and *Caenorhabditis briggsae*, which are included as outgroup species to root the phylogenetic tree. The subsequent analyses were done in two groups – those involving BUSCOs from the DpaA heterozygous region (6 orthologs – from DpaA, DpaB, DcoA, DcoB, *C. elegans* and *C. briggsae*), and those involving BUSCOs from the DpaA homozygous region (5 orthologs – from DpaB, DcoA, DcoB, *C. elegans* and *C. briggsae*).

For each BUSCO gene, our script called Muscle^94^ (v 5.1.0) to align the protein orthologs from each haploid genome. The script then turned the protein multiple sequence alignments into the corresponding cDNA multiple sequence alignments, after which any gaps in the alignments were eliminated by trimAl^95^ and the alignments were concatenated.

These concatenated alignments preserved the reading frames of the sequences. Concatenated alignments from both the DpaA heterozygous and homozygous regions were evaluated using modeltest-ng^32^ for the best nucleotide substitution models, with the nucleotides partitioned based on its position in the codon (1^st^, 2^nd^ or 3^rd^). For the BUSCOs found in the DpaA homozygous region, modeltest-ng results indicated that the first codon position should be evaluated using GTR+I, the second position using GTR+G4 and the third using GTR+I+G4. For the BUSCOs found in the DpaA heterozygous region, modeltest-ng results indicated that the first codon position should be evaluated using GTR+I, while the second and the third codon positions should be evaluated using GTR+I+G4. Finally, we constructed bootstrapped phylogenetic trees for the DpaA homozygous region and the Dpa heterozygous region using RAxML-ng^31^ and the evolutionary models determined by modeltest-ng above. Individual branches with poor bootstrap support (57% on one branch in the heterozygous tree, 53% on one branch in the homozygous tree) were collapsed into polytomic nodes. Trees were drawn with FigTree (http://tree.bio.ed.ac.uk/software/figtree/) using the .support output from RAxML (phylogenetic tree file containing bootstrap support values).

### Subtelomeric and telomeric array search

We used a Python script called TeloSearchLR^50^ to visualize the distribution of tandem repeats found at both ends of all the genomic reads. Natural DNA ends in a long-read sequencing library contain telomeres, which should be stranded: C-rich strand at the 5’ end, reading into the centre, and G-rich strand at the 3’ end, reading away from the centre of the chromosome. Repetitive sequences at sheared DNA ends tend not to have this bias. With this principle, we searched for 4-to 20-mer motifs (-k 4 –K 20) that are both enriched (-m 1 –M 100) and stranded at the read ends (-n 1000, capturing the information at the last 1000 bp of reads). The sequence TAAGGGTAAGGC was the most frequent stranded repeat motif found at the ends of sequencing reads. While TAAGGG also had stranded occupancy at the read ends, it did not appear as frequently.

Subtelomeric repeats were identified from chromosome-to-chromosome nucmer^24^ alignments, and manually annotated. Schematics for subtelomeric repeats were plotted using the package gggenes^96^ in R.

### Telomere length estimates using long-read sequencing libraries

We developed a Python script to estimate the lengths of the double-stranded portions of telomeres from a variety of nematode species. We identified several long-read genomic libraries derived from *C. elegans* and *C. briggsae* in addition to the *D. pachys* and *D. coronatus* libraries generated from this study (*C. elegans* strain VC2010/PD1074^97^ – SRR7594465, *C. elegans* strain CB4856^98^ – SRR8599835 to SRR8599843, *C. briggsae* strain QX1410^44^ – SRR17074503), chosen because their DNA isolation protocols did not have an excessive DNA shearing step, thus preserving the integrity of their telomeres.

We identified a set of probable telomeric sequencing reads by the strand-specific and terminal locations of repeated TTAGGC (*Caenorhabditis*) or TAAGGGTAAGGC (*Diploscapter*) as reported by TideHunter (v 1.4.2)^99^. Within this set of probable telomeric sequencing reads, we estimated the telomere lengths using two methods (**Supplementary fig. 7a**): 1) by finding the position of the last subtelomeric nucleotide (the “subtelomere mapping” method) and 2) by patterns of telomere repeat occupancy (the “repeat occupancy” method). For method 1 (“subtelomere mapping”), we aligned the reads to the known subtelomeric sequences from every chromosome to identify the position of the subtelomere-telomere boundary. We excluded the consideration of any telomeres that remain unresolved in the current assemblies – these are *C. elegans* CB4856 IIIL and VR; and *C. briggsae* QX1410 IL, IR, IVL, IVR, VL XL, and XR. For method 2 (“repeat occupancy”), we determined the occupancy of the telomeric repeat in two adjacent but non-overlapping 100-bp windows; the position between these two windows where the difference between these two occupancies is the greatest is the subtelomere-telomere boundary.

For all species, we tried to reconcile the telomere lengths estimated from these two methods. The error-prone nature of long reads meant that the subtelomere-telomere boundary estimated by method 2 often did not coincide perfectly with method 1 (**Supplementary Figure 7b**). In the sequencing libraries, we observed telomeric rearrangements that also led to discrepancies between the two estimates (**Supplementary Figure 7c**). Specifically in *Diploscapter*, we further observed that the subtelomere-telomere boundaries were often not the same, which meant that method 1 was less reliable (**Supplementary Figure 7d**). Accounting for all of this, for all telomere length estimates, we report the telomere length values using method 2 if and only if this estimate is within 50 bp of the estimated using method 1 in *Caenorhabditis*, or within 1000 bp using method 1 in *Diploscapter*.

### Fluorescent in-situ hybridization (FISH) of *D. pachys* repetitive subtelomeres

To generate FISH probes, a 2.8-kb fragment of the *D. pachys* subtelomere repeating unit was first cloned into pUC19. ∼1 µg of a purified PCR product amplified from the cloned subtelomere fragment was used to template the synthesis of digoxigenin (DIG)-labelled probes using DIG-High Prime (Roche 11745832910, Basel, Switzerland) following the manufacturer’s protocol.

*D. pachys* animals were cut open with 27G needles in a small volume of 2× egg buffer (50 mM HEPES pH 7.3, 1 mM EGTA pH 8, 4 mM EDTA pH 8, 236 mM NaCl, and 96 mM KCl) to extrude the gonads on a HistoBond microscope slide (VWR VistaVision 16004-406, Radnor, PA). An equivalent volume of 2× egg buffer with 8% formaldehyde was added to the dissection for a final fixative concentration of 4% formaldehyde. *D. pachys* gonad tissues were fixed for 5 minutes at room temperature before being flash-frozen on dry ice in the small flat space between a cover slip and the slide. The cover slip was removed using a razor blade while the slide was still frozen, and the frozen fixed tissue was washed once in ice-cold methanol and once in ice-cold acetone.

Hybridization of probes and the subsequent visualization were performed using a protocol originally developed by Dernburg and Sedat^100^. Briefly, the fixed tissues were rehydrated by submerging the slides for 10-15 minutes in a series of 2×SSCT buffers (0.2 M NaCl, 30 mM sodium citrate, 0.1% Tween-20) with increasing formamide concentrations (v/v): 0%, 20%, 40%, 50%, 50%. The hybridization mix was 10 µL of 0.3 M NaCl, 45 mM sodium citrate, 0.1% Tween-20, 50% (v/v) formamide, 50% (w/v) dextran sulphate and 1/1000 (v/v) of the probe DIG-labelling reaction, added directly to the rehydrated tissues with a coverslip on top. The coverslip was sealed on the slide with Elmer’s rubber cement (Westerville, OH) before the slide was heated to 95°C for 3 minutes to denature the genomic DNA. The slide was then slowly cooled to 37°C to allow the probes to hybridize overnight. With the coverslip off, probes were then washed away by submerging the slides 3 times in wash buffer (2×SSCT with 50% v/v formamide) at 37°C for 20 minutes each, and once each in wash buffers with decreasing concentrations of formamide at room temperature for 5-10 minutes each: 25%, 0%, 0%.

We used anti-digoxigenin antibodies to detect hybridized probes. First, the tissue was soaked in a blocking buffer (2×SSCT with 0.5% w/v bovine serum albumin [Sigma A3294]). Primary antibody staining was performed using a 1:500 dilution of the 21H8 monoclonal anti-DIG (Novus NBP2-31191 lot 20K30) in 2× SSCT, and secondary antibody staining was performed using a 1:400 dilution of the anti-mouse-IgG with FITC (Jackson 115-095-146) in 2× SSCT. The primary antibody staining was performed at 4°C overnight, while the secondary antibody staining was performed at room temperature for 2 hours, and each staining was followed by 3 washes in 2×SSCT. Fluorescent immunocomplexes were protected using a buffered anti-fade mounting solution with the nuclei counterstained with DAPI (Vectashield H-1200-10). Finally, images of the stained *D. pachys* gonads were captured on a Zeiss LSM880 AiryScan microscope using a 63× objective.

## DATA AVAILABILITY

Genomic sequencing reads from *D. pachys* and *D. coronatus* are available on the Sequence Read Archive (NCBI BioProject PRJNA1210125, PRJEB75588), where the *D. coronatus* assembled genome can also be found (GCA_964036155.1). The *D. pachys* genome has been submitted to NCBI, awaiting the assignment of an accession number. It is currently available via https://drive.google.com/open?id=1RE5PhDCd-YR2xh3r0kPEDxThM2R6FMVx&usp=drive_fs

## CODE AVAILABILITY

Algorithms for the analyses of the *Diploscapter* genomes, as well as code for generating figures in this work are deposited in the GitHub repository “GunsalusPiano/Diploscapter” (https://github.com/GunsalusPiano/Diploscapter).

## ACKNOWLEDGEMENTS

This work was supported by a gift from NYU Abu Dhabi (to FP), by NIH NIA grant R21 AG073830 (to KCG), by NSF BIO IOS 1656736 and NIH NIGMS grant R01GM141395 (to DHAF), and by the Wellcome Trust Grants 220540/Z/20/A and 218328 awards to the Wellcome Sanger Institute. We also thank Drs. Kohta Yoshida and Ralf Sommers for the HiC data from *Pristionchus exspectatus*^27^. For the purpose of Open Access, the author has applied a CC BY public copyright licence to any Author Accepted Manuscript version arising from this submission. We thank Philipp Schiffer for providing *D. corornatus* strain PDL0010.

## FIGURE TEXT

**Supplementary fig. 1: Transcriptional status of the telomerase protein component ortholog (*trt-1*) in *D. coronatus* and *D. pachys***.

Shown here are chromosomal coordinates (top), RNAseq reads mapping to the *trt-1* ortholog on a *Diploscapter* chromosome, and the *trt-1* ortholog gene model based on the RNAseq reads. **a**, the *trt-1* orthologs located on DcoA and DcoB are both actively transcribed. **b,** the *trt-1* orthologs located on DpaA and DpaB are transcribed, although transcripts from the first few exons are missing. For each ortholog, the t1 transcript is based on *ab initio* predictions as well as RNAseq data, while t2 (or t3) transcript is based solely on RNAseq data (see **Methods**).

**Supplementary fig. 2: Chromosome centre and arm characteristics for *P. exspectatus*, showing typical Rhabditid chromosome arm-centre differentiation**

a, Nigon painting of the *Pristionchus exspectatus* chromosomes, with 500-kb bins. **b**, Repeat content of *P. exspectatus* chromosomes along their lengths, in 100-kb bins. **c**, Exonic DNA content of *P. exspectatus* chromosomes along their lengths, 100-kb bins. **d**, GC content of *P. exspectatus* chromosomes along their lengths, 100-kb bins. **e**, Principal component 1 of *P. exspectatus* HiC data reveals regions of long-range intra– and inter-chromosome contacts. For all traits, two ancestral chromosome centres can be inferred from on the current *P. exspectatus* chromosome X. Data kindly provided by Yoshida & al^28^.

**Supplementary fig.3: Several traits within HFCs are markedly different from the rest of the genome**.

The chromosome-wide traits in Fig. 4 are plotted based on their locations – outside of HFCs vs. inside of HFCs for DpaA, DpaB, DcoA and DcoB. **a,** The long-range chromosomal interactions are not due to a higher density of restriction enzyme sites. In fact, HFC regions have lower restriction site density. **b,** The repeat densities outside and inside HFCs. *Diploscapter* HFCs have significantly fewer repeats. **c,** The exonic DNA densities outside and inside HFCs. *Diploscapter* HFCs are richer in exonic DNA. **d,** The GC content outside and inside HFCs. GC content is not statistically different outside vs. inside *Diploscapter* HFCs. *P*-values are calculated using Mann-Whitney *U* test.

**Supplementary fig. 4: *Diploscapter* orthologs of the Cel-TEBP contain a truncated POT-binding region (PBR)**.

**a,** A comparison of the domain structures TEBP-1/2 orthologs from *D. pachys, D. coronatus,* and *C. elegans*. The *C. elegans* TEBP-1 and TEBP-2 are known to bind *C. elegans* OB-fold proteins POT-1 and POT-2^36,37^. **b,** An alignment by Clustal Omega^58^ reveals that the *Diploscapter* orthologs of TEBP-1/2 are missing a large number of residues corresponding to the PBR (POT-binding region).

**Supplementary fig. 5: Long-term telomere dysfunction underlies karyotypic changes in *Diploscapter.***

**a**, The OB-fold protein ortholog is lost in the *Diploscapter* lineage, leading to changes in telomere length as telomerase may not be efficiently recruited to telomere ends. **b**, Telomeres shorten in this lineage and **c**, they are occasionally lost. **d**, Chromosome ends without telomeres are repaired incorrectly and are joined together. **e**, Chromosome ends are joined together whenever telomeric sequences are lost, until the entire haploid genome is fused into a single chromosome. Further fusions are not viable due to aneuploidy. **f**, Counteracting further telomere loss, ALT copies a ∼3-kb sequence to the chromosome ends. **g**, The ∼3-kb sequences diverge over time, and thus each chromosome end has a slightly different ∼3-kb sequence. **h**, Occasional telomere loss reactivates ALT, duplicating the 3-kb sequences at each end of the chromosome. Current *Diploscapter* chromosomes bear the molecular signature of this: short telomeres with long and repetitive subtelomeres

**Supplementary fig. 6: Two methods to estimate telomere lengths using error-prone long sequencing reads**

**a**, Telomere lengths can be estimated from reads by two methods. Method 1 uses the known subtelomeric sequences to determine the subtelomere-telomere boundary. Method 2 uses the change in the occupancy of the telomeric repeat motif in two adjacent non-overlapping windows, where *a* or *b* = fraction occupancy of the telomeric motif in their respective window. Where the difference between *a* and *b* is the largest (*max*[*b – a*]) is the subtelomere-telomere boundary. Estimates by these two methods can differ due to the phenomena described below. **b**, sequencing errors in the telomere can lead to a discrepancy between the estimates from the two methods. **c**, Rearrangements near the telomere can also lead to a discrepancy between the two estimates. **d**, In *Diploscapter*, the subtelomere-telomere boundary can vary, leading to yet another type of discrepancy between the two telomere estimation methods. A telomere length is only considered when these two estimates are reasonably close (see **Methods**).

