## Supplementary figure 1 for "Genome evolution in parthenogenetic nematodes shaped by chromosome rearrangements and rare sex"

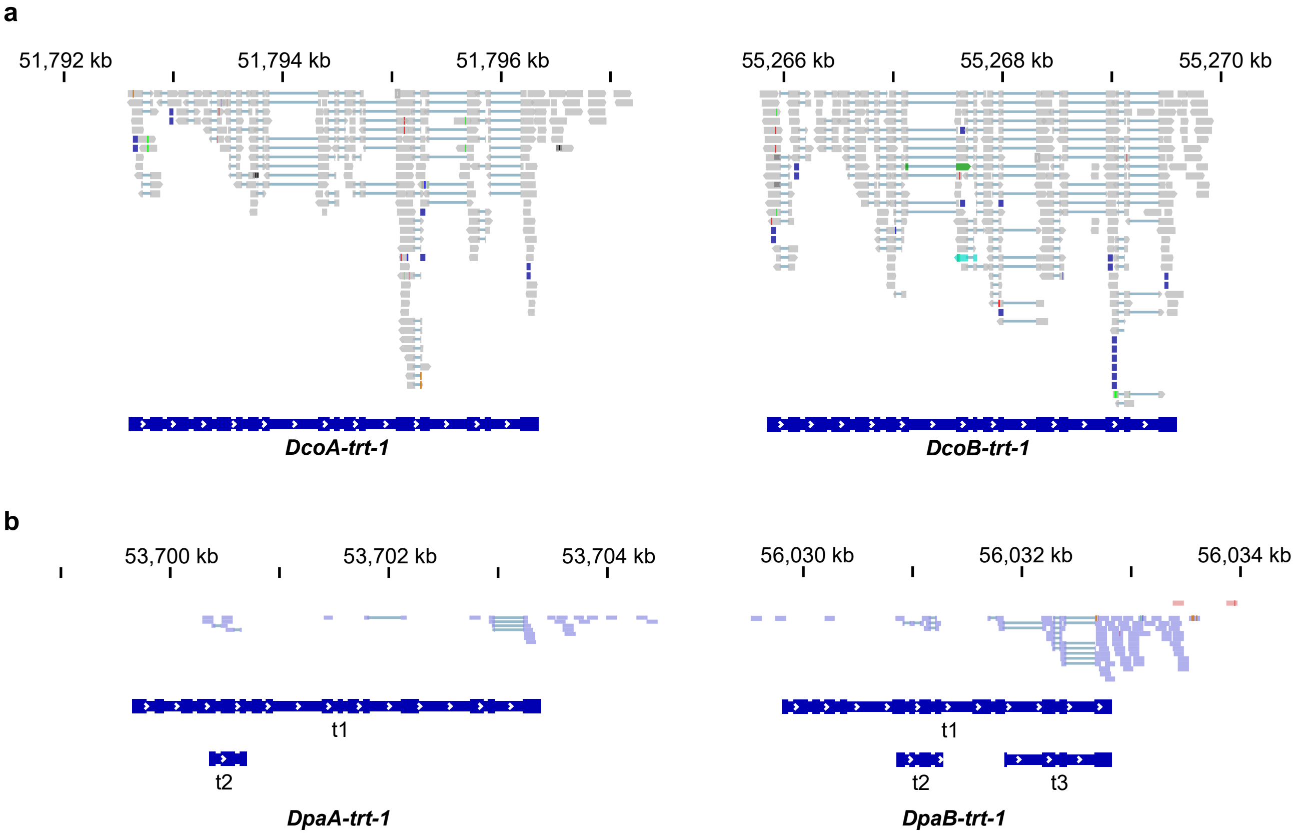


**Supplementary fig. 1: Transcriptional status of the telomerase protein component ortholog (*trt-1*) in *D. coronatus* and *D. pachys*.**
Shown here are chromosomal coordinates (top), RNAseq reads mapping to the *trt-1* ortholog on a *Diploscapter* chromosome, and the *trt-1* ortholog gene model based on the RNAseq reads. **a**, the *trt-1* orthologs located on DcoA and DcoB are both actively transcribed. **b,** the *trt-1* orthologs located on DpaA and DpaB are transcribed, although transcripts from the first few exons are missing. For each ortholog, the t1 transcript is based on *ab initio* predictions as well as RNAseq data, while t2 (or t3) transcript is based solely on RNAseq data (see **Methods**).
