## Supplementary figure 2 for "Genome evolution in parthenogenetic nematodes shaped by chromosome rearrangements and rare sex"

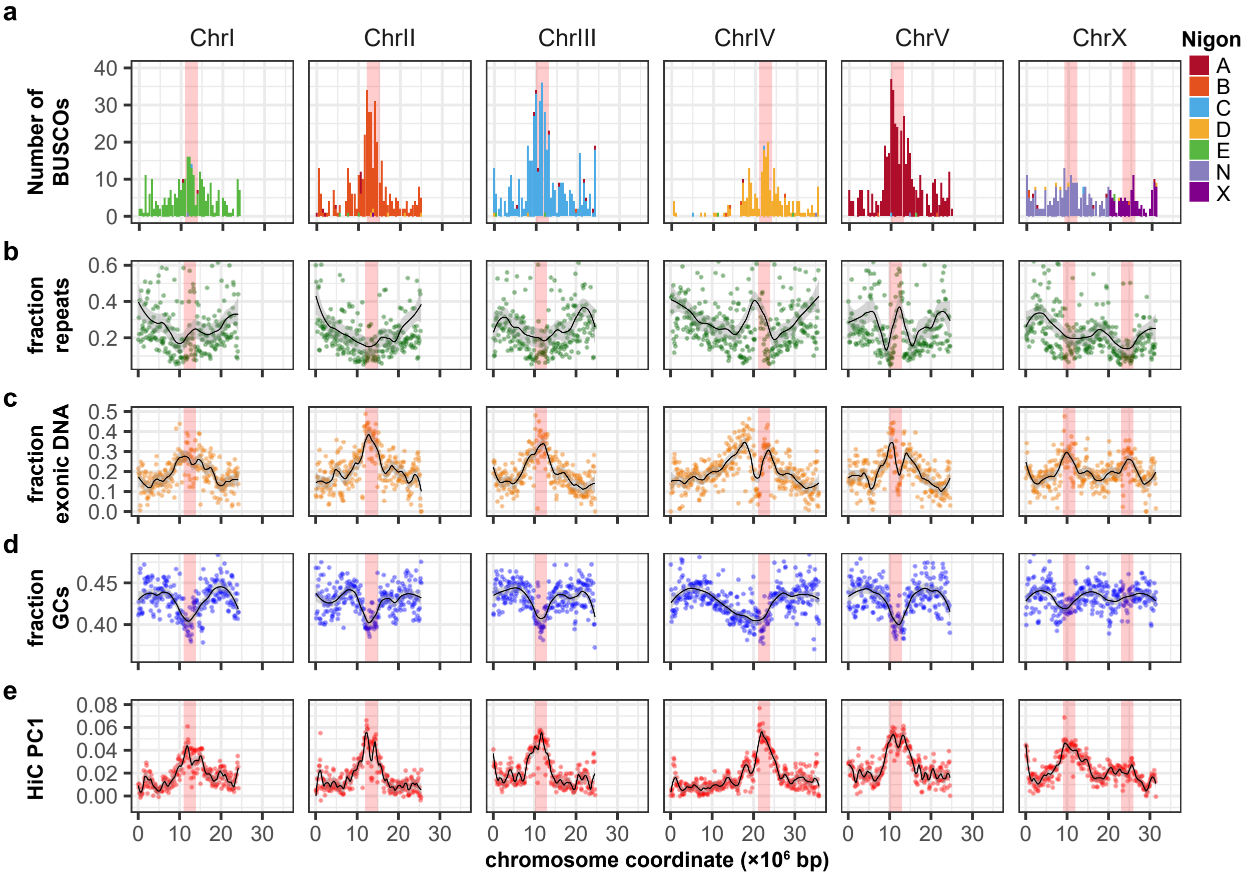


**Supplementary fig. 2: Chromosome centre and arm characteristics for *P. exspectatus*, showing typical Rhabditid chromosome arm-centre differentiation**

**a**, Nigon painting of the *Pristionchus exspectatus* chromosomes, with 500-kb bins. **b**, Repeat content of *P. exspectatus* chromosomes along their lengths, in 100-kb bins. **c**, Exonic DNA content of *P. exspectatus* chromosomes along their lengths, 100-kb bins. **d**, GC content of *P. exspectatus* chromosomes along their lengths, 100-kb bins. **e**, Principal component 1 of *P. exspectatus* HiC data reveals regions of long-range intra- and inter-chromosome contacts. For all traits, two ancestral chromosome centres can be inferred from on the current *P. exspectatus* chromosome X. Data kindly provided by Yoshida & al[28].
