## Supplementary figure 3 for "Genome evolution in parthenogenetic nematodes shaped by chromosome rearrangements and rare sex"

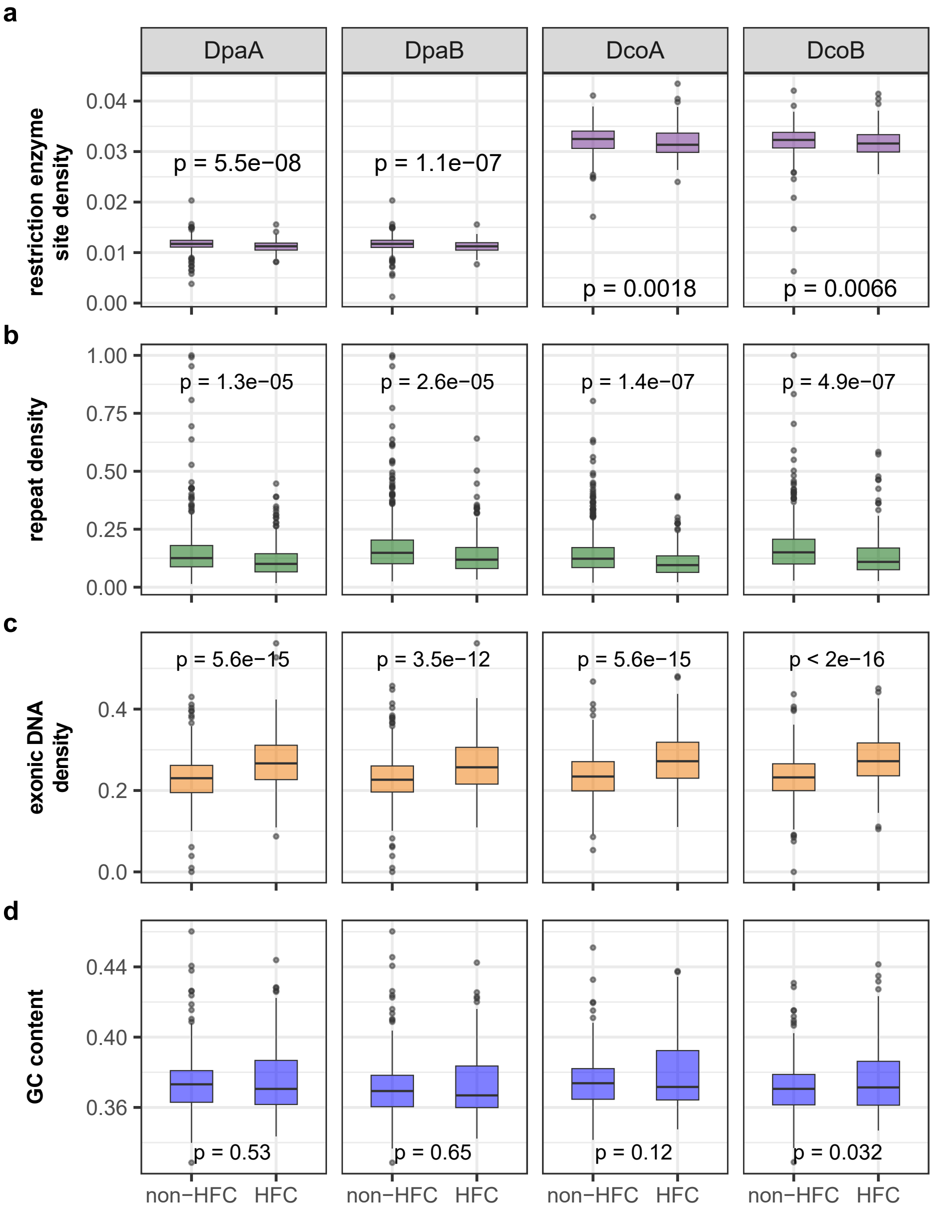


**Supplementary fig.3: Several traits within HFCs are markedly different from the rest of the genome.**
The chromosome-wide traits in Fig. 4 are plotted based on their locations - outside of HFCs vs. inside of HFCs for DpaA, DpaB, DcoA and DcoB. **a,** The long-range chromosomal interactions are not due to a higher density of restriction enzyme sites. In fact, HFC regions have lower restriction site density. **b,** The repeat densities outside and inside HFCs. *Diploscapter* HFCs have significantly fewer repeats. **c,** The exonic DNA densities outside and inside HFCs. *Diploscapter* HFCs are richer in exonic DNA. **d,** The GC content outside and inside HFCs. GC content is not statistically different outside vs. inside *Diploscapter* HFCs. *P*-values are calculated using Mann-Whitney *U* test.
