## Supplementary figures and images for "Genome evolution in parthenogenetic nematodes shaped by chromosome rearrangements and rare sex"

### Supplementary figure 4

#### Supplementary fig. 4

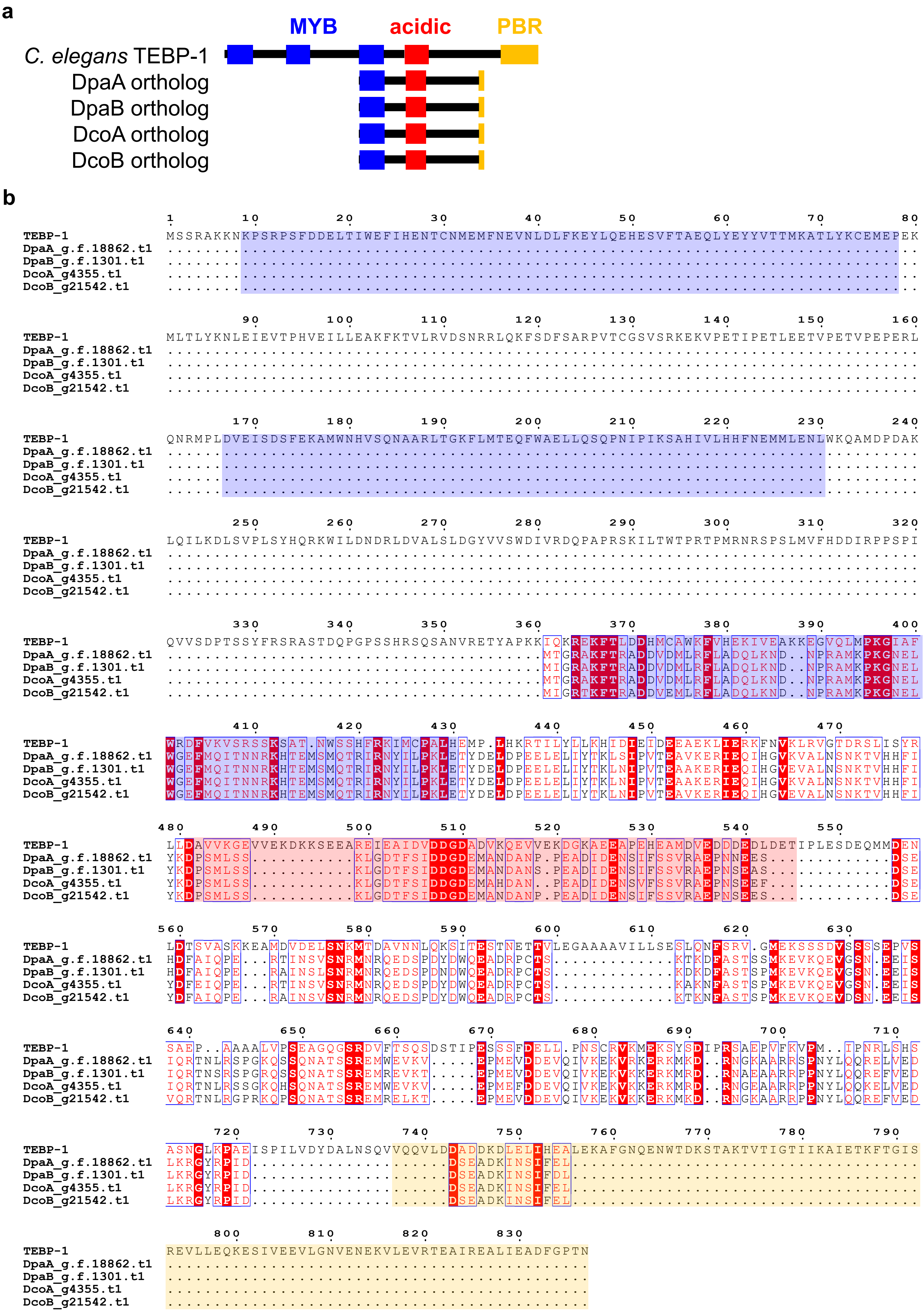
