## Supplementary figure 5 for "Genome evolution in parthenogenetic nematodes shaped by chromosome rearrangements and rare sex"

**Supplementary fig. 5**

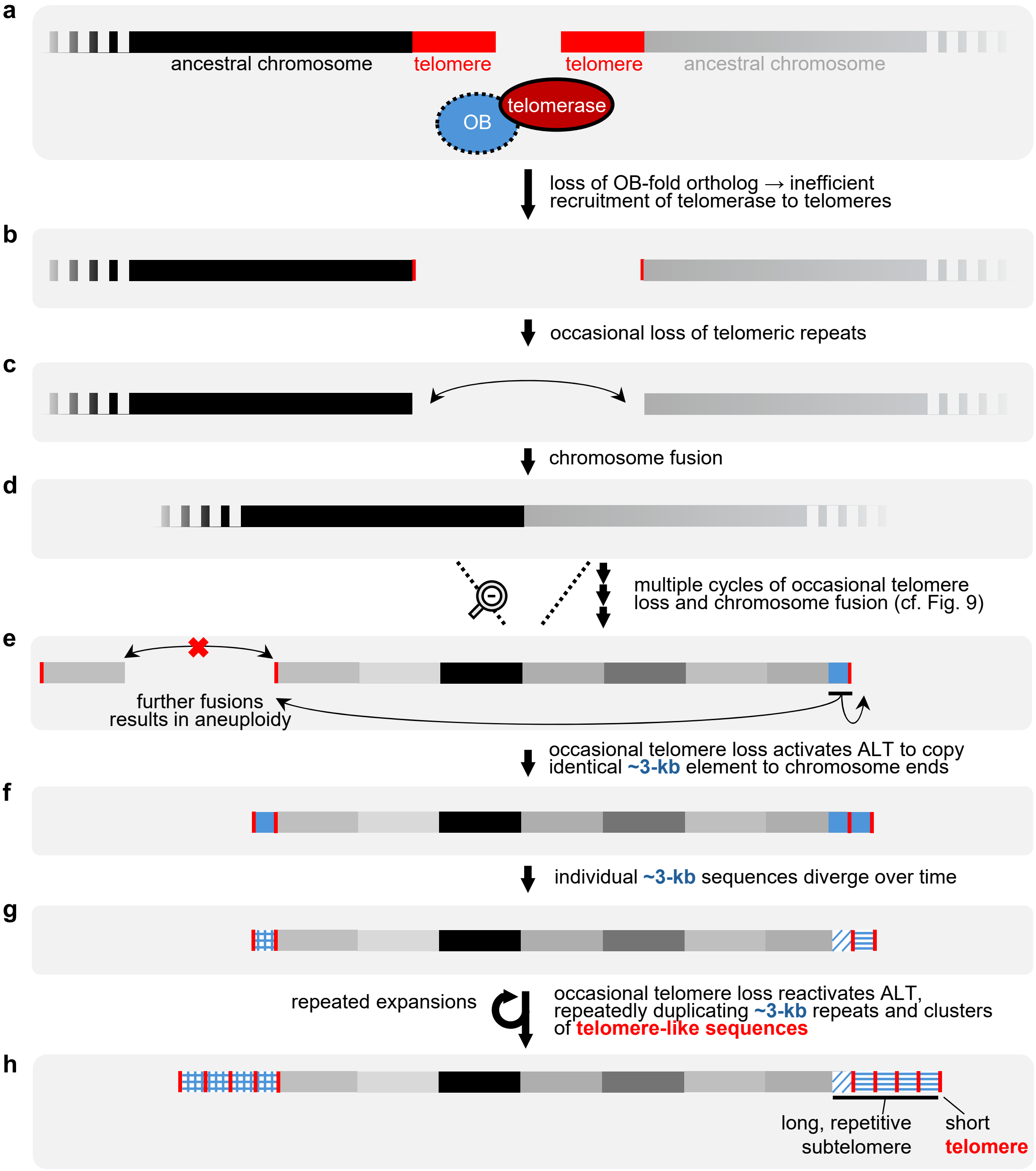


**Supplementary fig. 5: Long-term telomere dysfunction underlies karyotypic changes in *Diploscapter.***

**a**, The OB-fold protein ortholog is lost in the *Diploscapter* lineage, leading to changes in telomere length as telomerase may not be efficiently recruited to telomere ends. **b**, Telomeres shorten in this lineage and **c**, they are occasionally lost. **d**, Chromosome ends without telomeres are repaired incorrectly and are joined together. **e**, Chromosome ends are joined together whenever telomeric sequences are lost, until the entire haploid genome is fused into a single chromosome. Further fusions are not viable due to aneuploidy. **f**, Counteracting further telomere loss, ALT copies a ~3-kb sequence to the chromosome ends. **g**, The ~3-kb sequences diverge over time, and thus each chromosome end has a slightly different ~3-kb sequence. **h**, Occasional telomere loss reactivates ALT, duplicating the 3-kb sequences at each end of the chromosome. Current *Diploscapter* chromosomes bear the molecular signature of this: short telomeres with long and repetitive subtelomeres
