## Supplementary figure 6 for "Genome evolution in parthenogenetic nematodes shaped by chromosome rearrangements and rare sex"

**Supplementary figure 7**

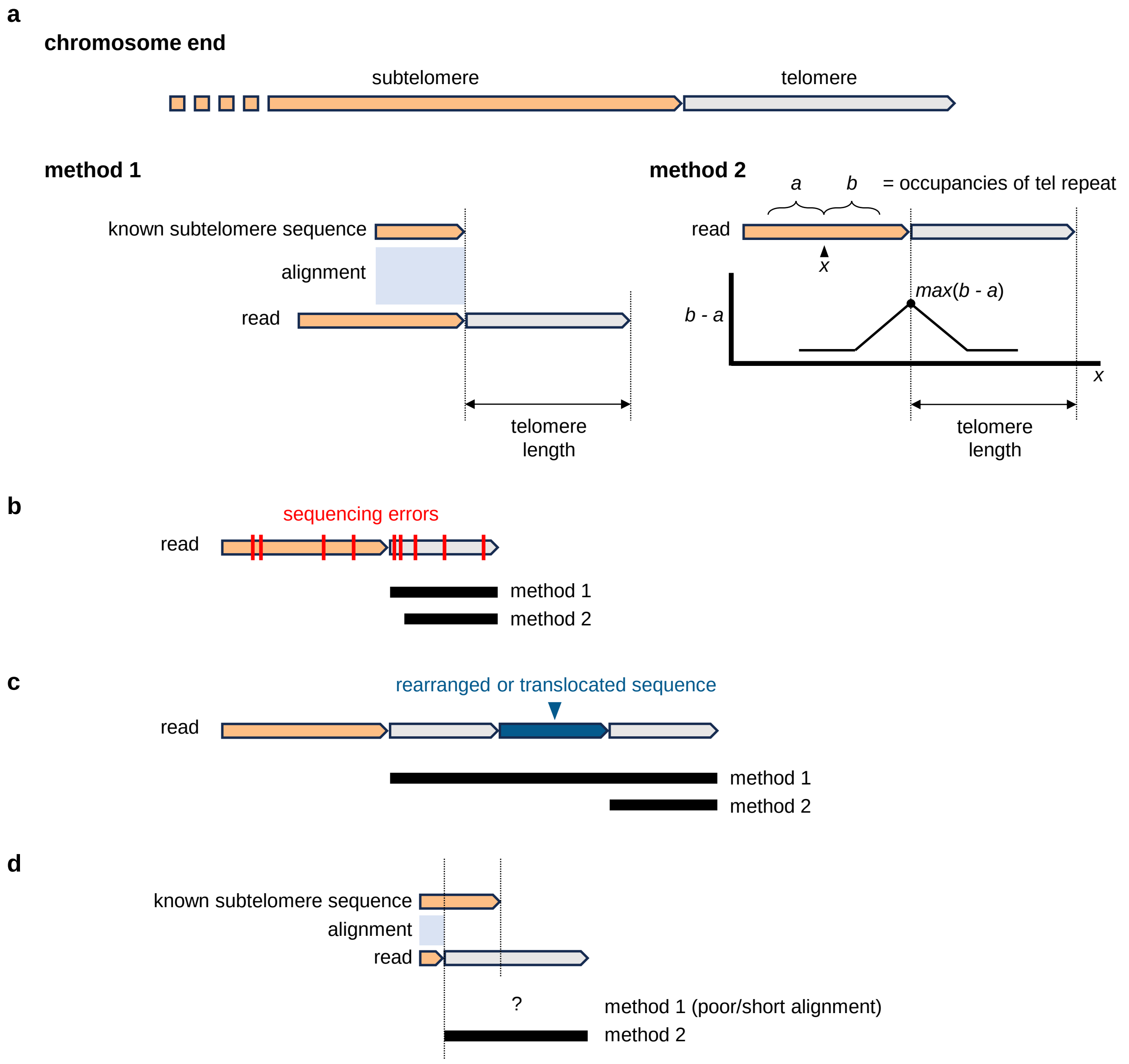


**Supplementary fig. 7: Two methods to estimate telomere lengths using error-prone long sequencing reads**

**a**, Telomere lengths can be estimated from reads by two methods. Method 1 uses the known subtelomeric sequences to determine the subtelomere-telomere boundary. Method 2 uses the change in the occupancy of the telomeric repeat motif in two adjacent non-overlapping windows, where *a* or *b* = fraction occupancy of the telomeric motif in their respective window. Where the difference between *a* and *b* is the largest (*max*[*b* - *a*]) is the subtelomere-telomere boundary. Estimates by these two methods can differ due to the phenomena described below. **b**, sequencing errors in the telomere can lead to a discrepancy between the estimates from the two methods. **c**, Rearrangements near the telomere can also lead to a discrepancy between the two estimates. **d**, In *Diploscapter*, the subtelomere-telomere boundary can vary, leading to yet another type of discrepancy between the two telomere estimation methods. A telomere length is only considered when these two estimates are reasonably close (see **Methods**).
