## Supplementary table 1 for "Genome evolution in parthenogenetic nematodes shaped by chromosome rearrangements and rare sex"

| Species | *Diploscapter pachys* | *Diploscapter coronatus* |
|---|---|---|
| Sequencing technology | Oxford Nanopore R9.4.1 | PacBio Sequel IIe |
| Assembler | Canu, het settings | hifiasm |
| Scaffolding technique | PoreC | HiC |
| Scaffolder | YaHS[74] | YaHS[74] |
| Number of scaffolds | 2 nuclear,1 mtDNA | 2 nuclear, 1 mtDNA |
| Assembly size (bp) | 155,979,874* | 169,703,327 |
| Assembly N50 (bp) | 85,103,350 (chromosomal) | 87,757,044 (chromosomal) |
| G+C content | 0.372 | 0.373 |
| Number of protein-coding genes | 31,963* | 34,641 |

**Table 1: Key statistics of the two *Diploscapter* assemblies.**
For the *D. pachys* assembly, an asterisk (*) indicates a value determined for the assembly where the homozygous region is represented only once.
