## Supplementary table 2 for "Genome evolution in parthenogenetic nematodes shaped by chromosome rearrangements and rare sex"

| chromosome end | nominal length of the subtelomere (bp) in the assembly | subtelomere repeat motif length (bp) | number of repeat copies in the assembly | estimated subtelomere length (bp) | estimated total number of copies |
|---|---|---|---|---|---|
| DpaA-L or DpaB-L* | 39,362 | 2,807 | 14.0 | 39,362 | 14.0 |
| DpaA-R | 68,594 | 3,200 | 21.4 | 756,500 | 236.4 |
| DpaB-R | 44,662 | 3,008 | 14.8 | 92,641 | 30.8 |
| DcoA-L | 25,068 | 2,879 | 8.7 | 94,771 | 32.9 |
| DcoA-R | 16,264 | 3,547 | 4.6 | 87,135 | 24.6 |
| DcoB-L | 14,531 | 3,201 | 4.5 | 93,944 | 29.3 |
| DcoB-R | 23,185 | 3,249 | 7.1 | 53,248 | 16.4 |

**Supplementary table 2: Estimated subtelomere lengths in *D. pachys* and *D. coronatus*.** The fold excess read depth in the subtelomeres over the rest of the chromosome allows the estimation of subtelomere lengths. *The homozygous left end of DpaA and DpaB appear to be fully resolved in our assembly, complete with telomeric array of TAAGG[G/C] repeats.
